# A single *Microcoleus* species causes benthic cyanotoxic blooms worldwide

**DOI:** 10.1101/2023.10.13.562201

**Authors:** Pilar Junier, Guillaume Cailleau, Mathilda Fatton, Pauline Udriet, Isha Hashmi, Danae Bregnard, Andrea Corona-Ramirez, Eva di Francesco, Thierry Kuhn, Naïma Mangia, Sami Zhioua, Daniel Hunkeler, Saskia Bindschedler, Simon Sieber, Diego Gonzalez

**Author notes:** co-corresponding authors: Pilar Junier, Rue Emile-Argand 11, CH-2000, Neuchatel. Diego Gonzalez, Rue Emile-Argand 11, CH-2000, Neuchatel. Simon Sieber, Winterthurerstrasse 190, CH-8057, Zurich.

## Abstract

Recently, proliferations of benthic cyanobacteria producing derivatives of anatoxin-a have been reported in rivers all over the world. In three river systems, in New Zealand, the USA, and Canada, a cohesive cluster of *Microcoleus* strains was responsible for toxin production. Here, we document a similar toxigenic event that occurred at the mouth of the river Areuse in lake Neuchâtel (Switzerland) and caused the death of several dogs. Using 16S RNA-based community analysis, we show that riverine benthic communities are dominated by Oscillatoriales and especially by *Microcoleus* strains. We correlate the detection of one sequence variant with the presence of anatoxin-a derivatives and use metagenomics to assemble a complete circular genome of the strain. The strain is distinct from the ones isolated in New Zealand, the USA, and Canada, but belongs to the same species; it shares significant traits with them, in particular a relatively small genome and incomplete vitamin biosynthetic pathways. Overall, our results suggest that the major anatoxin-a-associated benthic proliferations worldwide can be traced back to a single ubiquitous species, *Microcoleus anatoxicus,* rather than to a diversity of cyanobacterial lineages. We recommend that this species be monitored internationally in order to help predict and mitigate similar cyanotoxic events.

## Introduction

Cyanobacterial blooms pose a significant threat to aquatic ecosystems and human health. They can interfere with the survival of both plants and animals by sequestering nutrients and causing oxygen depletion. Many cyanobacteria also produce toxic metabolites, collectively called cyanotoxins, which are active against a wide range of potential competitors and predators, from bacteria to vertebrates ^1–6^. Commonly reported cyanotoxins in aquatic environments include microcystins, cylindrospermopsin, anatoxin-a congeners, and saxitoxins ^7^. Because of their potency and wide target-range, these toxins can by themselves profoundly affect the functioning of ecosystems ^8^. They also represent a direct danger for human health during recreational activities and could have far-reaching and long-term consequences since some of them have been shown to bioaccumulate along the trophic chain ^9^.

While a majority of reported blooms are caused by cyanobacteria proliferating in the water column, blooms of substrate-attached, i.e. benthic, species are increasingly documented ^10,11^. This rise in benthic blooms is likely linked to human activities and, possibly, to global warming, which makes their investigation a pressing task ^12,13^. Benthic blooms typically start with the thickening and widening of bacterial mats on the bed of rivers and lakes ^13^. At some point, the mats, loaded with oxygen entrapped in the filaments, detach from the substrate and reach the surface of the water bodies. Benthic blooms are associated with many different toxins, but most commonly with anatoxin-a ^10^. Anatoxin-a (ATX) and its congeners, homoanatoxin-a (HTX), dihydroanatoxin-a (dhATX), and dihydrohomoanatoxin-a (dhHTX), are potent agonists of the nicotinic acetylcholine receptors ^14^. They interfere with the function of neuromuscular junctions and can cause respiratory arrest in dogs and other vertebrates. ATX-producing benthic blooms have been reported from all over the world and particularly well documented in New Zealand (Hutt river, but many others), in the USA (Russian and Eel rivers), and more recently in Canada (Wolastoq river) ^10,14–21^.

In New Zealand, the USA, and Canada, ATX-producing benthic blooms are caused by strains belonging to the *Microcoleus* genus (Oscillatioriales, section III) ^22^. Due to the uncertainties of cyanobacterial taxonomy, these strains were initially classified as *Tychonema, Oscillatoria,* or *Phormidium,* but are progressively being renamed *Microcoleus* since a taxonomic clarification proposed in 2013 ^23^. Among the ATX congeners, dhATX is the most commonly found in *Microcoleus*-dominated benthic blooms in New Zealand and the USA, while ATX seems the most frequent in Canada ^19–21,24^. Blooms typically comprise more than one strain with the overall toxin concentration correlating with the relative proportion of toxigenic to nontoxigenic strains ^16,25,26^. Proliferation positively correlates with dissolved inorganic nitrogen-concentrations and in some studies with temperature and light intensity; lower phosphate concentrations also seem to favor blooms, possibly owing to the capacity of the implicated strains to efficiently mobilize phosphate from the substrate ^13,27,28^.

Partial, fragmented genomes have been assembled for several ATX-producing strains isolated in New Zealand, the USA, and Canada ^20–22^. All these strains encode a specific variant of the ATX-biosynthesis cluster first described in *Cylindrospermum stagnale.* It comprises a short *anaG* allele leading to ATX, rather than HTX, as the synthesis intermediate. In addition, the presence of *anaK*, which encodes for an oxidase, is responsible for the transformation of ATX into dhATX ^29,30^. This is consistent with the detection of dhATX as the major ATX congener in these strains in New Zealand and the USA; some of the Canadian strains have *anaK* controlled by a different promoter that may be less active. Strains from New Zealand, the USA, and Canada are closely related at the nucleotide level and have common genomic features compared to other *Microcoleus* strains, in particular a smaller genome size and missing genes in common vitamin biosynthetic pathways ^22^. Among them, only one strain has been properly described, *Microcoleus anatoxicus* Stancheva & Conklin, whose type strain is PTRS2 from the Russian river in the USA ^20^. How the sequenced *Microcoleus* strains from New Zealand and Canada relate to the *M. anatoxicus* species is not entirely clear and the overall genetic diversity of ATX-congener synthesizing *Microcoleus* strains is unknown.

Here, we document a case of ATX-producing benthic cyanobacterial bloom, which happened in summer 2020 at the mouth of the Areuse in the Lake of Neuchâtel (Switzerland). This event entailed the death of six dogs and was followed by similar events in several other rivers and lakes in Switzerland. We use 16S rRNA gene sequencing and metagenomics to identify the strains responsible for the toxin production and to survey the diversity of benthic cyanobacterial species in the river. While ATX-associated blooms and death of dogs have been reported in many rivers and lakes in Europe, this is the first such event that is fully documented at the genomic level.

## Results

### A cyanotoxic event associated with the proliferation of a Microcoleus strain

By the end of July 2020, six dogs died after being in contact with cyanobacterial scum close to the mouth of the river Areuse at the shore of the Lake Neuchâtel (Switzerland). The toxicological analyses performed on the dogs identified anatoxin-a (ATX), a widespread toxin from cyanobacterial origin, as the most likely cause of death. The scum was particularly dense at the mouth of the Areuse, most likely because it originated from the river bed and got amassed in the delta by the backflow of the lake. The scum was composed of mats of entangled filaments often enclosing gas bubbles which increased their buoyancy, an appearance typical of benthic cyanobacterial blooms ^10,13^.

To single-out the microorganisms responsible for the dogs’ poisoning, we sampled the floating mats accumulating at the mouth of the river right after the event (July 31st) and two weeks later (August 16th) (Figure 1AB). Under the light microscope, the mats were mostly composed of filamentous cyanobacteria, greenish to dark-brown in color. Their trichomes presented a smooth morphology common to many benthic Oscillatoriales (Figure 1CD). Single cells were about 5-10 µm in width and square in shape, with hardly any constriction between individual cells and without sign of differentiation (no akinetes or heterocysts); at the tip of the filament, cells could be rounded or conical, and often bore calyptra. We narrowed down the taxonomic identification by sequencing a partial 16S rRNA gene amplicon using cyanobacterial primers (STable 1). Consistent with the microscopy, the closest matches of the consensus sequence in 16S rRNA gene databases were Oscillatoriales belonging to the *Microcoleus* genus (sometimes under their basonymes *Phormidium*, *Tychonema,* or *Oscillatoria*). To determine the full bacterial composition of the scum, we used high-throughput 16S rRNA gene amplicon (V3-V4 region) sequencing on the initial samples taken after the incident (end of July; 7cyano2) and a sample taken on the same site two weeks after (mid-August; 7cyano1). Floating mats were largely dominated by one *Microcoleus* amplicon sequence variant (ASV) representing more than 65% of the bacterial reads in both samples (Figure 1EFG). Flavobacteria and Proteobacteria (*Rhodobacter, Rhodoferax, Brevundimonas*) were the most abundant heterotrophs in these samples (Figure 1EF). Overall, our observations and analyses suggested that the cyanotoxic event in the lake Neuchâtel was caused by the proliferation of at least one *Microcoleus* strain capable of producing an anatoxin-a congener.

**Figure 1.**
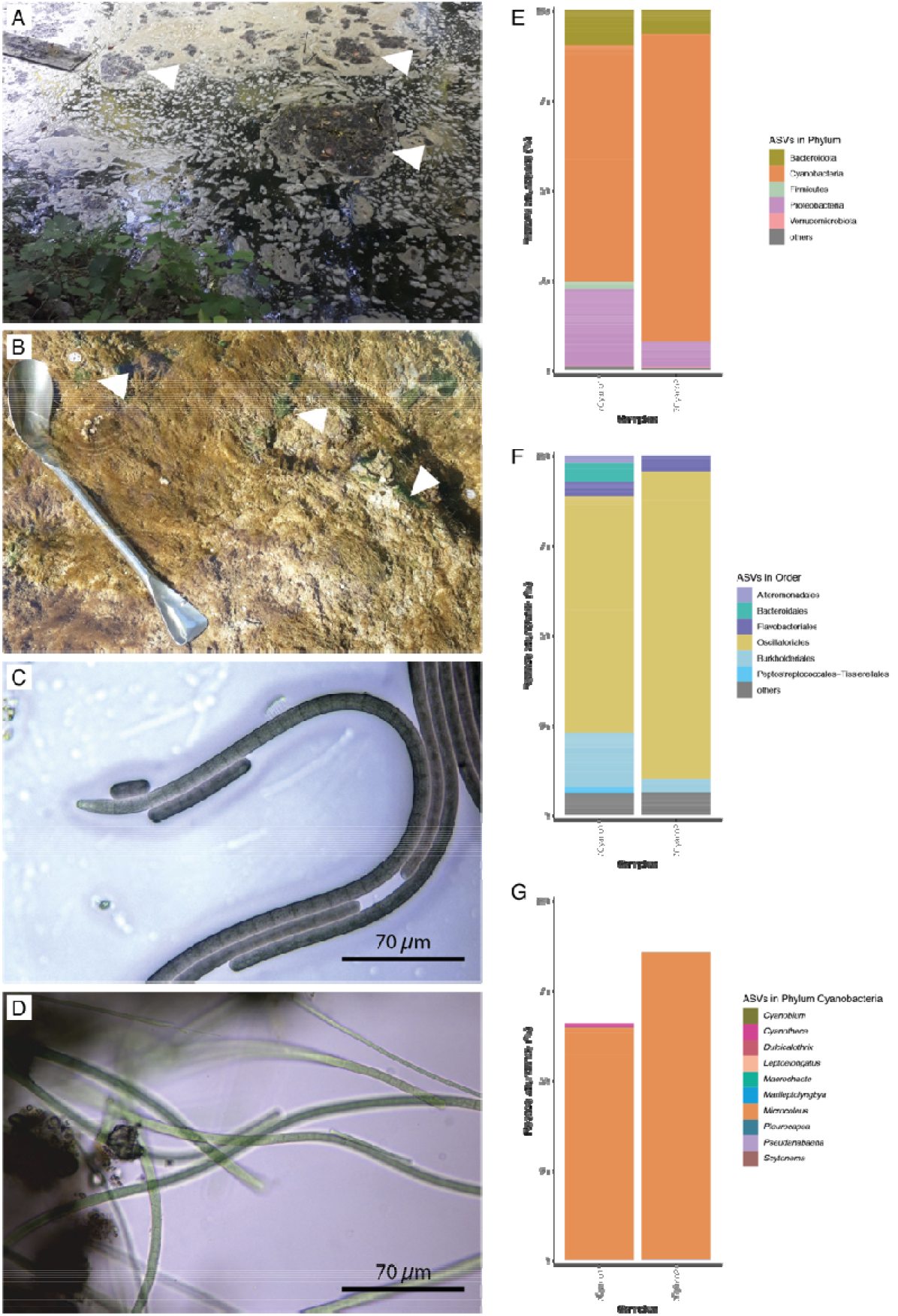
The initial toxigenic bloom event (July-August 2020). A. Mats and scum accumulating at the surface of the river close to the mouth of the Areuse. The low level of the river and a backflow of Lake Neuchatel created an area of low flow and accumulation of biomass in which scums (floating mats) were clearly observed. Arrows indicate floating mats among the scum. B. Benthic mats on stones with bright green patches (arrows) dominated by *Potamolinea* sp. C. Trichomes composing the toxigenic mats collected close to the mouth of the Areuse (dominant species. *Microcoleus* anatoxicus). These trichomes lack branching, heterocysts, and akinetes, and are characteristic of Oscillatoriales. Phase contrast microscopy, objective 100x. D. Trichomes composing the bright green mats collected close to the mouth of the Areuse (dominant species: *Potamolinea* sp.). Phase contrast microscopy, objective 100x. E. and F. The composition of the bacterial communities in the floating mats collected close to the mouth of the Areuse at Phylum (E) and the Order (F) levels. The communities present in two samples were analyzed on the basis of the V3V4 region of the 16S rRNA gene. G. The cyanobacterial fraction of the bacterial community is clearly dominated by *Microcoleus* at the genus level.

### Geographic origin, distribution, and diversity of blooming strains

We next addressed the geographic origin and distribution of the blooming strains along the river Areuse. In early September, we took multiple samples of cyanobacterial mats or biological material growing on immerged rocks, wood or sediments, at three main locations: in the delta on the river bed (11 samples, both mats and sediments; STable 1), on the banks along the last 1 km of the river (23 samples, both mats and sediments; STable 1), and at a dam located five kilometers upstream of the mouth, close to the village of Noiraigue (7 samples, floating mats only; STable 1). Floating mats were only observed near the mouth of the river and in Noiraigue. At the mouth of the river, the bed was almost entirely covered in dark microbial mats that resembled the floating scums (SVideo 1); these appeared to accumulate mainly in the delta of the river and were mostly absent on the lake floor (SVideo 2). A distinct type of floating cyanobacterial mats, whose color was bright green, was also collected in the lower part of the Areuse (samples 34, X, Y1 and Y2). While samples collected near the delta appeared clearly dominated by *Microcoleus*-like cyanobacteria under the microscope, the rock surface samples on the river banks contained a diversity of photosynthetic organisms, including cyanobacteria, algae, and diatoms.

To get a sense of the diversity and frequency of filamentous cyanobacteria and, especially, of *Microcoleus* strains in the different samples, we performed high-throughput 16S rRNA gene amplicon sequencing (V3-V4 region) on extracted DNA. Filamentous cyanobacteria were largely dominant in scums and floating mats, representing up to 85% of all bacterial reads (Figure 2AB); they were very well represented in most other samples (sediments) with at least 20% of identified reads (with the exception of samples 100 and 200 from the upper Areuse). Among cyanobacteria, two operational taxonomic units could reach very high frequencies in a subset of samples: *Microcoleus*, present in most samples, and *Potamolinea,* dominating the four samples containing bright green filaments (samples 34, X, Y1, Y2). At the ASV level, the *Microcoleus* genus included a significant diversity with at least 20 different ASVs present across samples. One of these ASVs, identified as *Microcoleus anatoxicus* ^20^, was dominant in the scum and floating mats sampled in the delta and in the upper Areuse (Noiraigue). *Microcoleus* / *Tychonema* ASVs were more diverse in other samples especially in the mid-river bed. Like in the initial sampling, the most frequent heterotrophs across samples belonged to the Proteobacteria and Sphingobacteria (Flavobacteria) phyla.

**Figure 2.**
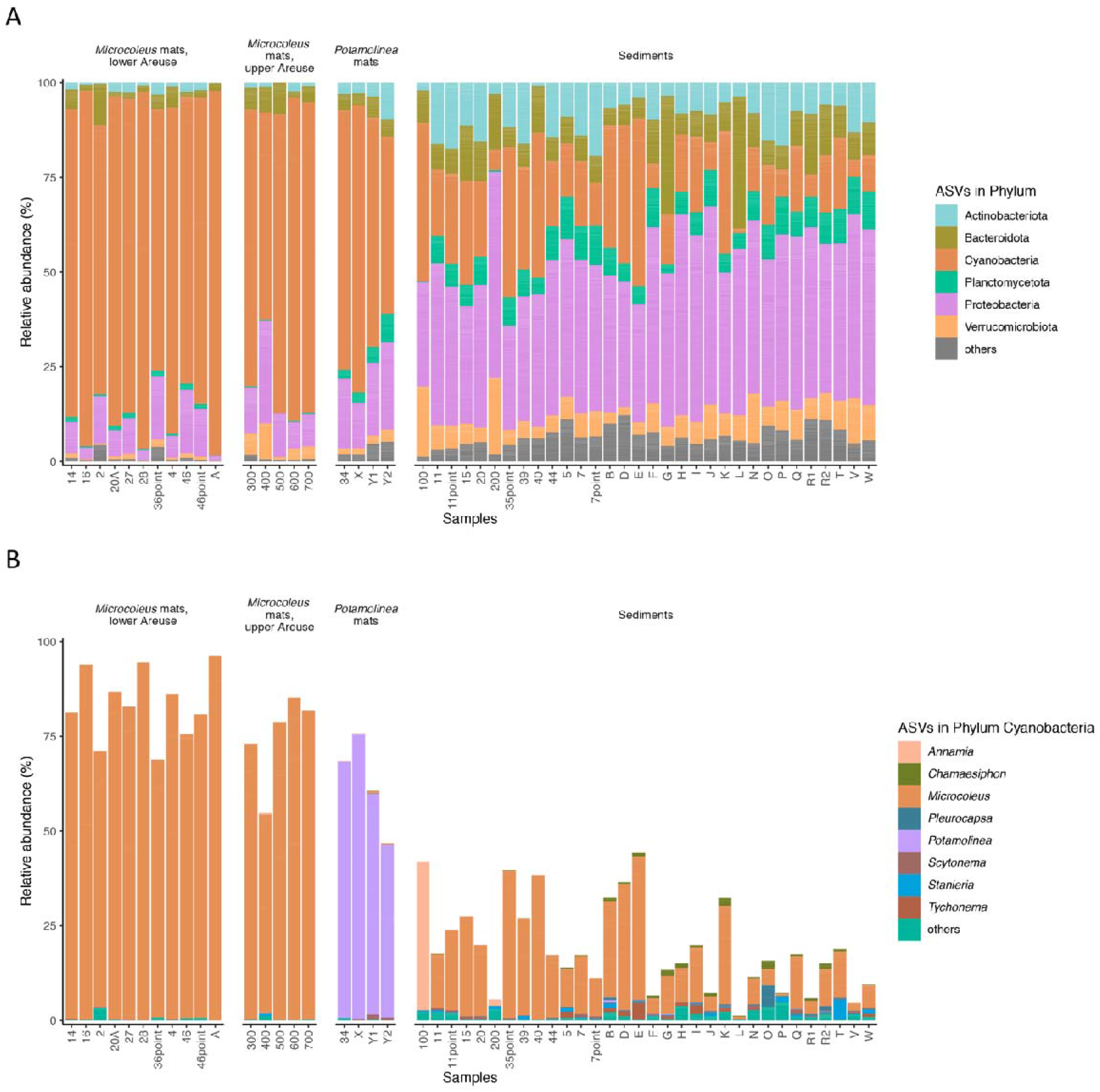
Composition of the bacterial communities present in mats and sediments from the Areuse in September 2020. The community composition is based on the V3V4 region of the 16S rRNA gene. The samples have been arranged by type (cyanobacterial mats *vs* sediments), dominant species (*Microcoleus vs Potamolinea*) and, for mats, by location (Lower Areuse *vs* Upper Areuse / Noiraigue). A. Community analysis at the phylum level, showing a substantial presence of Cyanobacteria and Proteobacteria in collected mats and sediments. B. Community analysis at the genus level, showing the dominance of *Microcoleus* in the cyanobacterial fraction of most samples.

Overall, these results indicate that *Microcoleus* strains are abundant in benthic mats in both their floating (blooming) and their surface-attached forms. One ASV in particular is associated with blooms in the lower and the upper Areuse.

### A single Microcoleus ASV is associated with a genetic marker for anatoxin-a and higher anatoxin-a and dihydro-anatoxin-a concentrations

In parallel to the community profiling, we used published primers targeting the *anaC* gene to test each sample for the presence of the ATX biosynthesis cluster ^31^. While virtually all the samples from the delta of the Areuse were positive for the *anaC* marker, the same marker was only detected in a quarter of the samples along the lower part of the river and in floating mats in the upper part of the Areuse (STable 1). This suggests that while the ATX producing strain is sporadically present on the bed and banks of the Areuse all along the river, it only reaches densities high enough to be detected by PCR in the delta region where the river current meets the backflow of the lake.

The presence of ATX and dhATX was confirmed by analyzing the samples by Ultra-high-performance liquid chromatography (UHPLC) coupled with a high-resolution mass spectrometer (HR-MS) ^32^ or a triple quadrupole mass spectrometer (TQMS) ^33^. The extract from floating mats dominated by *Microcoleus* contained 0.1 % and 5% of ATX and dhATX respectively (11 µg of ATX-a and 241 µg of dhATX in a 8.2 mg of extract). In contrast, the samples dominated by *Potamolinea* contained less than 0.04% of ATX and dhATX (9.5 µg of ATX in 35.7 mg of extract).

All samples that tested positive for *anaC* contained a significant proportion of one *Microcoleus* ASV. While there was a clear association between that ASV and the ATX genetic marker, genomic data were still needed to formally demonstrate that the identified strain was the one actually encoding the ATX biosynthesis cluster.

### The anatoxin-a-producing strain from the Areuse is closely related to Microcoleus anatoxicus PTRS2 and to other Microcoleus strains from New Zealand, the USA, and Canada

To formally identify the ATX-producing *Microcoleus* strain, we used a long-read high-throughput sequencing technology (Pacific Biosciences SMRT) to sequence DNA extracted from four samples. Three samples (A and N from the lower Areuse and 600 from the upper Areuse) were selected because they presented a high relative abundance of the *Microcoleus* ASV of interest and were positive for the *anaC* marker; the fourth sample (X from the lower Areuse) was chosen because it was dominated by the other blooming strain, *Potamolinea*. For all four samples, we obtained a number of large contigs that were binned based on coverage and tetranucleotide identity, and classified based on global homology and full 16S rRNA gene sequences (SFigures 1-4, STable 2). We were able to assemble the full circular genome of a *Microcoleus* strain for sample A and N, as well as large contigs of a *Microcoleus* strain for sample 600 and large contigs of a *Potamolinea* strain for sample X. As expected, the contigs assembled from sample X (*Potamolinea* sp.) belonged to a different branch of the cyanobacterial phylogenetic tree, clustering with Coleofasciculates close to Nostocales (SFigure 5).

The full *Microcoleus* genomes assembled from samples A and N (strain NeuA/N) were virtually identical (>99.99% average nucleotide identity [ANI]) (Figure 3B). They significantly differed from the fragmentary genome assembled from sample 600 (Neu600) based on ANI (<90%) (Figure 3A), compositional binning (SFigure 1), and full 16S rRNA gene sequence (26 nucleotide differences over the whole sequence, although the closest match for both is *Microcoleus anatoxicus* strain PTRS2). Phylogenetically, all three strains belonged to the *Microcoleus*-*Tychonema* branch of Oscillatoriales, but several strains from public databases branched between strain NeuA/N and Neu600. The NeuA/N strain formed a monophyletic clade with the dhATX-producing strains from the USA and New Zealand (Figure 3A, SFigure 5) and showed high conservation over the 16S rRNA gene (>98% identity; SFigure 6). Average nucleotide identity with strains from the USA, New Zealand, and Canada was in general above 95%, the species threshold (Figure 3B, SFigure 7) ^34^. Based on digital DNA-DNA hybridization, strain NeuA has a probability higher than 50% to belong to the same species as representative dhATX/ATX producing strains from New Zealand, the USA, and Canada, including the taxonomic reference *Microcoleus anatoxicus* strain PTRS2 (estimated DNA-DNA hybridization between 60% and 70%, STable 3) ^35,36^. Consistent with the whole genome comparisons, the secondary structure of the D1-D1’ helix of the 16S-23S rRNA genes internal transcribed spacer (ITS) is identical in *Microcoleus* sp. NeuA, *Microcoleus* sp. EPA2, and *Microcoleus anatoxicus* strain PTRS2 (SFigure 8). Overall, this strongly suggests that the dhATX/ATX producing *Microcoleus* strains proliferating in rivers of temperate countries worldwide belong to the same species.

**Figure 3.**
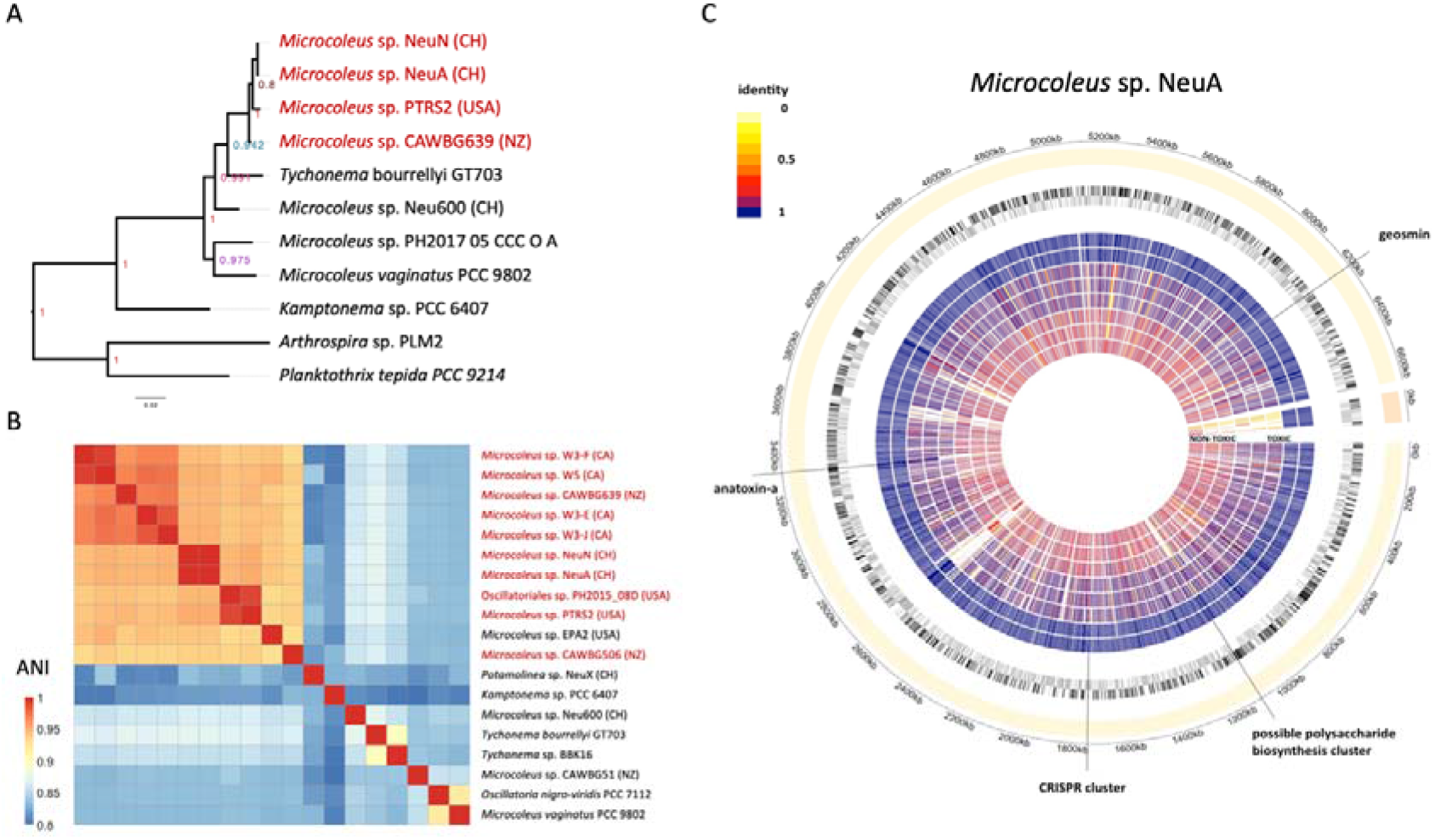
Phylogenomic characteristics of *Microcoleus* strains isolated from the Areuse. A. Phylogenetic tree of strains closely related to *Microcoleus* sp. NeuA based on 28 conserved proteins. The tree shows that dhATX-synthesizing strains from New-Zealand, the USA, and Switzerland belong to the same clade; *Microcoleus* sp. Neu600 branches before the acquisition of the dhATX biosynthesis cluster. B. Average nucleotide identity (ANI) among strains closely related to *Microcoleus* sp. NeuA. The figure indicates that NeuA/N (Switzerland), CAWBG639 (New Zealand), PTRS2 (USA), and four strains from the Wolastoq river (Canada) likely belong to the same species (ANI>=0.94). Both phylogeny and ANI suggest *Tychonema bourrellyi* GT703 and *Microcoleus* sp. Neu600 are the closest relatives of the dhATX-cluster encoding clade. C. Circular plot of *Microcoleus* sp. NeuA genome. From the exterior to the interior: a. molecules (yellow: genome; orange: plasmid) and coordinates; b. open reading frames (black: + strand; grey: - strand); c. protein identity value between reference (*Microcoleus* sp. NeuA) and *Microcoleus* sp. NeuA (toxic), NeuN (toxic), Neu600 (toxic), PTRS1 (toxic), CAWBG639 (toxic), EPA2 (lost dhATX cluster), Tyc600 (nontoxic), CAWBG58 (nontoxic), PH2017_15_JOR_U_A (nontoxic), PCC 7112 (nontoxic).

### The genome content of Microcoleus sp. NeuA

The genome of *Microcoleus* sp. NeuA includes one circular chromosome of 6.5 MB and one large circular plasmid of 0.1 MB; it encodes 5’687 protein-coding genes, 72 tRNAs, and three rRNA operons (Figure 3C). At the protein level, the strain shares many features with the dhATX-producing strains from the USA and New Zealand. A number of broad features distinguishes them from closely related strains that do not encode the dhATX biosynthesis cluster, in particular the presence of a geosmin biosynthesis cluster, a set specific of CRISPR/Cas proteins, and a large cluster of genes of uncertain function including several polysaccharide biosynthesis or modification enzymes (Figure 3C).

*Microcoleus* sp. NeuA encodes a full ATX biosynthesis cluster comprising 10 genes (Figure 4A). This cluster includes a short *anaG* allele and the additional *anaK* gene characteristic of organisms that synthesize dhATX as their main toxin congener ^29^. The complement and order of the genes is similar to the one found in the dhATX-producing *Microcoleus* strains from New Zealand, the USA, and Canada with the exception of a variable region, between *anaB-G* and *anaK-I*, encoding hypothetical or uncharacterized proteins (Figure 4A). Overall, these results confirm that the dhATX-producing *Microcoleus* strains from New Zealand, the USA, Canada, and Switzerland (NeuA/N) have similar properties including in terms of toxin-synthesis capacity. By contrast, the contigs assembled from sample 600 did not include an ATX biosynthesis cluster, even fragmentary. A detailed examination of the flanking regions of the ATX biosynthesis cluster and their homologues in the non-toxic strains, including strain 600, clearly shows three different genomic states (Figure 4B): i) an ancestral state before the insertion of the ATX biosynthesis cluster, found in *Tychonema bourrellyi* FEM_GT703, *Tychonema* sp. BBK16, and our *Microcoleus* sp. Neu600; ii) a state where the full cluster is present at the same position, found in *Microcoleus* sp. CAWBG639, PTRS1 and NeuA/N; and iii) a third state found in *Microcoleus* sp. EPA2 (USA), where most of the cluster has been excised with a leftover of a hundred bases. The phylogenetic tree in Figure 4C shows the most likely history of the strain family, showing the acquisition of the ATX biosynthesis cluster at the beginning of the crown branch and its subsequent loss on the EPA2 branch.

**Figure 4.**
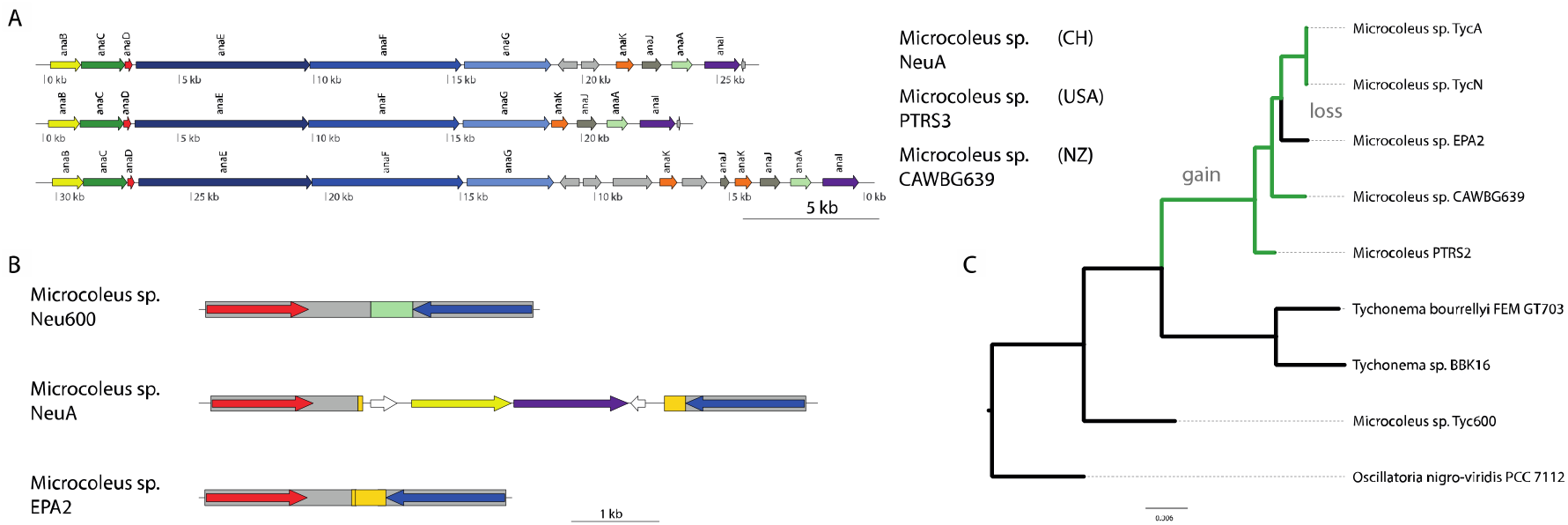
Sequence and evolution of the dhATX biosynthesis cluster in *Microcoleus* species. A. dhATX biosynthetic clusters from *Microcoleus* sp. NeuA, PTRS1 (USA), and CAWBG639 (NZ). The clusters are very conserved except for a variable region between the two parts of the cluster (*anaBCDEFG* and *anaKJAI*). B. The genomic context of the dhATX cluster in the ancestral nontoxic state (Neu600), after dhATX cluster acquisition (NeuA), and after secondary loss (EPA2). In green, a portion of DNA found in the ancestral state only; in orange, a portion of the dhATX biosynthesis cluster, which remains in EPA2 after cluster loss. C. Reconstruction of gain and loss events on a phylogenetic tree.

Besides the ATX cluster, the genome of *Microcoleus* sp. NeuA/N encodes three additional biosynthesis clusters that present homology with those of geosmin, anachelin (a cyanobacterial siderophore), and scytocyclamide (STable 4). The dhATX-synthesizing *Microcoleus* strains from New Zealand and the USA have been reported to differ from non-producing *Microcoleus* strains by several genomic traits, in particular a smaller genome size (<7MB) and the presence of incomplete or alternative metabolic pathways ^22^. The traits reported in these reference genomes hold true for the NeuA/N Swiss strain, but are not always exclusive to toxigenic strains in our dataset. Overall, we find that, compared to nontoxic strains, the dhATX producers tend to lack some genes in vitamin biosynthetic pathways, namely thiamine (*tenA* coding for thiaminase II), pyridoxal (*pdxH* coding for pyridoxamine 5’-phosphate oxidase), and cobalamine (*cbiG* coding for cobalt-precorrin 5A hydrolase), to have more genes involved in sugar metabolism (for instance, UDP-glucose epimerase *galE*) and polysaccharide modification (sugar-kinases and phosphatases), as well as lipid and lipopolysaccharide metabolism, and to have a different set of CRISPR-cas9 associated proteins (SFigure 9-11).

### Bacterial communities associated to different mat-forming cyanobacteria are distinct

Literature suggests that ATX-producing *Microcoleus* strains harbor specific cyanosphere-associated bacteria ^37^. To test whether our data supported this hypothesis, we did a principal component analysis at the species level on the mat datasets, excluding all cyanobacterial ASVs. Samples dominated by *Microcoleus* sp. NeuA, *Microcoleus* sp. Neu600, and *Potamolinea* sp. clustered separately, with three species belonging to the Sphingobacteriales—*Pedobacter* sp., *Paludibacter* sp., and *Flavobacterium* sp.—being the major drivers of the separation (Figure 5A). Consistent with this, a correlation analysis indicates that *Microcoleus* is positively correlated with the *Pedobacter* genus and negatively correlated with the *Potamolinea* genus (Figure 5B). A co-occurrence network analysis based on the lower Areuse samples confirms the association between *Microcoleus* sp. NeuA and *Pedobacter* as well as the negative relation between *Microcoleus* sp. NeuA and *Potamolinea* (Figure 5C). Overall, these results strongly suggest that the bacterial communities assembled around different mat-forming cyanobacteria present a different composition.

**Figure 5.**
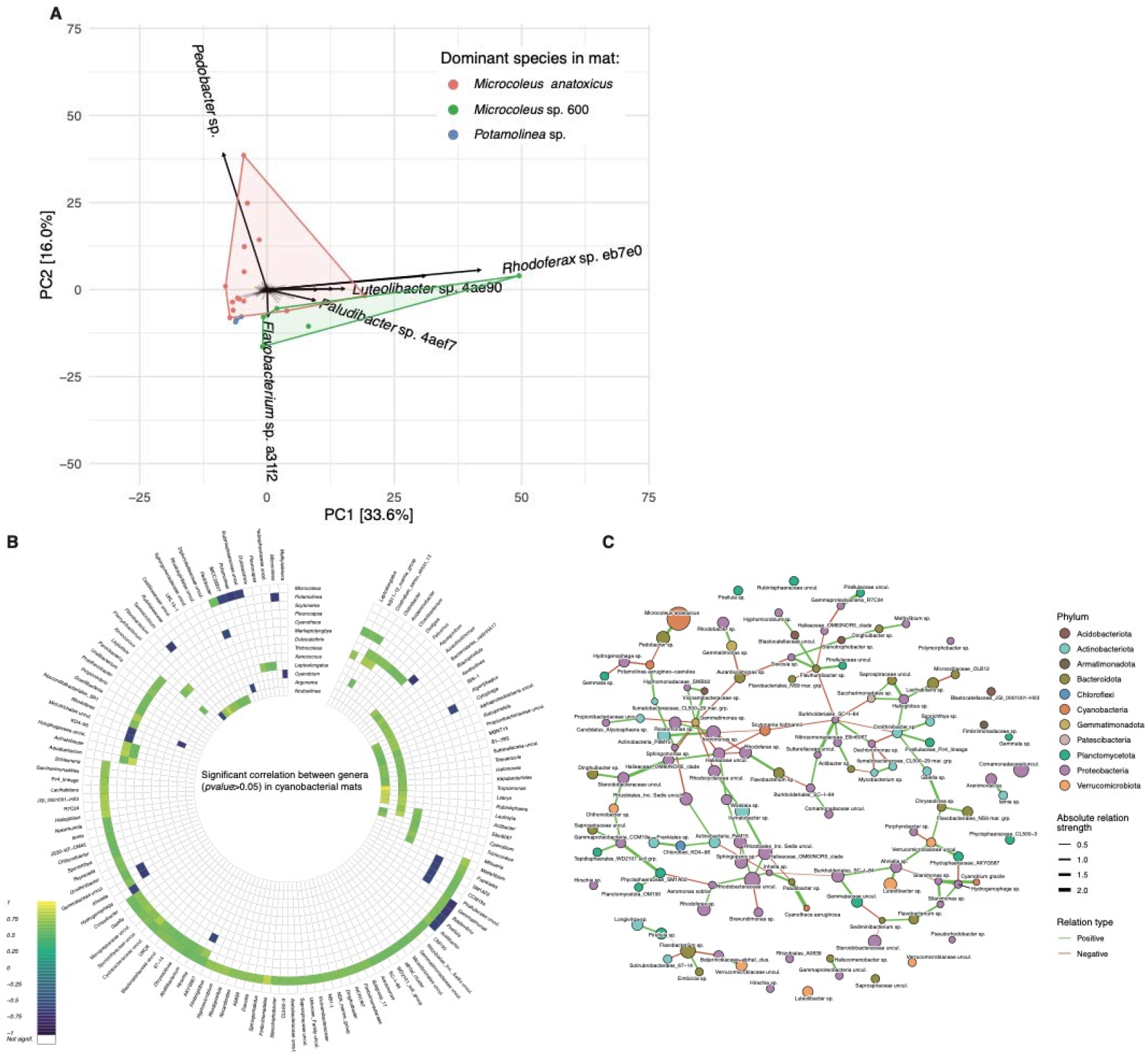
Bacterial communities in benthic mats differ in composition depending on the dominant cyanobacterial species. A. Principal component analysis (PCA) of bacterial communities in mat samples excluding all cyanobacterial ASVs (species level). All samples in the three first groups shown in Figure 2 are represented colored by the dominant cyanobacterium (*Microcoleus* sp. NeuA and sp. Neu600, *Potamolinea* sp.). The two first dimensions of the PCA are sufficient to separate the three groups of associated bacterial communities, which suggests that the communities are distinct from one another. B. Significant correlations (Spearman, p-value < 0.05) between cyanobacterial genera and all bacterial genera represented in more than two samples (mats only). C. Co-occurrence network reconstruction from the mat samples (excluding the upper Areuse mats), showing that the toxigenic *Microcoleus* species presents positive co-occurrence with *Pedobacter* sp. and negative co-occurrence with *Potamolinea* sp. Nodes are colored by Phylum; edge color indicates positive or negative co-occurrence; edge width represents co-occurrence strength.

## Discussion

The dog poisoning event that happened in 2020 by the lake Neuchâtel adds to an increasing list of episodes in which livestock, domestic animals, and even humans have been affected by the consumption of cyanotoxin-containing scums or mats ^38,39^. This event resulted from the expansion of toxigenic *Microcoleus* strains, producing high levels of dhATX and ATX. The toxin synthesis might have been stimulated upon exposure to light and oxygen at the surface of the lake and on the shores, since physiological stress is known to increase toxin levels by several orders of magnitude ^40^. In the case reported here, the compound that actually led to the death of the dogs was probably dhATX, whose concentration was about ten times higher than ATX in the mats (STable 1). Given the LD50 of dhATX (2.5 mg/kg by gavage), reported to be lower than ATX (10.6 mg/kg), the dogs would have had to ingest several grams of wet mats to reach the lethal dose ^41^.

While the 2020 event had exceptional spatial extension and consequences, the toxigenic species is likely well-established in the Areuse, given its presence at different densities in different types of samples taken several kilometers apart in the river (Figures 1 and 2). Moreover, once an habitat has been colonized by cyanobacteria, subsequent proliferations become more likely, which suggests that similar events may increase in frequency in the future in the Areuse ^10^. *Microcoleus*-dominated benthic proliferations in rivers tend to occur at intermediate flow rates, high nitrogen (>1mg/L total nitrogen) and low phosphate (<0.01 mg/L total phosphorus) concentrations ^28,42^. Such conditions are prevalent in the lower part of the Areuse based on our observations and official water quality data. The exceptional accumulation of mats close to the mouth of the river may have been prompted by low flow rates in the river and strong backflow coming from the lake, both preventing any flushing of accumulating mats during the weeks before the event ^38^.

Two toxins, dhATX and ATX, as well as a genetic marker of the ATX biosynthesis cluster, were mainly detected in samples dominated by a single amplicon sequence variant belonging to a *Microcoleus* strain. This is consistent with recent literature on toxin benthic blooms, which repeatedly identified *Microcoleus* strains, sometimes referred to as *Tychonema* sp. or *Phormidium* sp., as the main producers of ATX congener in rivers ^17,21,22,43–45^. This ASV was present in a majority of submerged samples and was clearly dominant in floating mats (delta, river banks and upper Areuse) except in the bright green blooms dominated by *Potamolinea* (Figure 1D) ^46^. This ASV (V3-V4 region) corresponds to the 16S rRNA gene sequence found in both *Microcoleus* sp. NeuA/N (dhATX producer) and *Microcoleus* sp. Neu600 (nonproducer), making them indistinguishable in the 16S rRNA gene community analysis. This highlights the co-occurrence of different closely related filamentous cyanobacterial strains within benthic mats, which suggests that competition for dominance might actually be frequent in benthic communities. Sample 600 from the upper Areuse is an interesting case: it was dominated by the nonproducer strain Neu600 (based on the long-read shotgun sequencing data), but tested positive for *anaC* and presented levels of toxin similar to samples dominated by strain NeuA/N. This suggests that a small proportion of an ATX producing strain, maybe a close relative of NeuA/N, was present in the sample and possibly overproduced the toxins. Such cases of mixed blooms of ATX-producers and nonproducers have been commonly observed in New Zealand ^26^. Whether interactions within such mats are cooperative—the toxic strain helping the nontoxic strain to proliferate, for instance by deterring crustaceans or worms—or competitive—the toxic strain slowly outcompeting the nontoxic one—is unknown. The presence of *Potamolinea* spp. also appears to limit the dominance of *Microcoleus* sp. and correlates with a lack of toxin detection. This may hint at potential competition between strains of the two genera. Future studies on the co-existence and/or competition between toxigenic and non-toxigenic *Microcoleus* species or between *Potamolinea* species and *Microcoleus* species may shed light on the population dynamics of benthic cyanobacteria.

Using high-throughput long-read sequencing of environmental samples, we successfully assembled a full circular genome for *Microcoleus* sp. NeuA/N (Figure 3). This strain is closely related to *Microcoleus anatoxicus* strain PTRS2, based on a conserved protein phylogeny, the 16S rRNA gene sequence (99.07% identity), the average nucleotide identity of aligned regions in the chromosome [ANI] (96.04%), digital DNA-DNA hybridization [dDDH] (61.6-67.4%, recommended method), and structure of the D1-D1’ helix of the ITS between the 16S and 23S rRNA genes (Figure 3, SFigure 5B and 8). Except for dDDH, which is slightly lower than the usually accepted 70% species threshold with the method used, all other criteria support the classification of the strain sequenced from the Areuse as *Microcoleus anatoxicus* strain NeuA/N. Moreover, we present evidence that different dhATX- or ATX-producing *Microcoleus* strains from New Zealand, the USA, and Canada (CAWBG639, CAWBG506, PTRS1-3, EPA2, and several strains from the Wolastoq river) likely belong to the same species. Although dDDH between strains is again somewhat lower than the usually accepted 70% threshold (STable 3), ANI and nucleotide identity of the 16S rRNA genes, whenever a full sequence is available, remain very high, i.e. >94% and >98% respectively (Figure 3B, SFigure 6). The high ANI, in particular, suggests that allele exchange is sufficient among these strains to keep recombination rates high and justify, from a biological viewpoint, to include them within a single species, specifically *M. anatoxicus.* However, we would need more full circular genomes of *M. anatoxicus* and closely related strains to precisely adjust the different species-delineation criteria and find meaningful thresholds for the *Microcoleus* genus.

The genome of *M. anatoxicus* strain NeuA/N encodes a variant of the ATX biosynthesis cluster known to generate dhATX as the main toxin congener, which is consistent with the toxicological results ^22,29^. While the organization of the locus is very similar to the ones found in the *Microcoleus* CAWBG639 and PTRS1-3 strains from New Zealand and the USA respectively, as well as in contigs assembled from the Wolastoq river in Canada, it is clearly distinct from the model cluster found in *C. stagnale* (Nostocales) (Figure 4) ^22,29^; it is also distinct from the clusters found in phylogenetically closer benthic Oscillatoriales like *Kamptonema* PCC 6506, which produces HTX as its main congener ^30^. This highlights the diversity of extant ATX biosynthesis clusters in Cyanobacteria and their potential to synthesize a variety of chemical derivatives of ATX, some of which might still be unidentified. Compared to other Oscillatoriales, the genome of *M. anatoxicus* strain NeuA/N is rather small with 6.5 MB plus a 100 kb plasmid. The size difference is partly due to a smaller number of secondary metabolite biosynthetic clusters and to a trend towards gene loss in conserved vitamin biosynthetic pathways (SFigures 9-11). These distinctive traits may be specific to the phylogenetic branch in which the ATX biosynthesis cluster got introduced or may represent subsequent adaptations to toxin production or growth in blooms.

*Microcoleus*/*Phormidium*-associated bacterial communities are thought to be quite similar across countries at higher taxonomic levels ^47^. In our dataset, however, the bacterial communities assembled around benthic cyanobacteria appear to be species-specific at lower taxonomic levels (genus and species) (Figure 5): closely related species like *M. anatoxicus* strain NeuA/N and *Microcoleus* sp. 600, for instance, harbor clearly distinct communities (Figure 5A). We find that Sphingobacteriales (especially *Pedobacter* in our case) may play a central role in the economy of communities surrounding *M. anatoxicus*. Future studies should address whether the potential vitamin auxotrophies of the dhATX-producing *Microcoleus* strains may be balanced in nature by an association with specific strains, especially Sphingobacteriales, that may provide the missing compounds.

In conclusion, this study shows a clear convergence between cyanotoxic events happening at remote sites on the planet. These events are caused by strains belonging to a cohesive cluster around the *M. anatoxicus* type strain PTRS2 and encoding a biosynthesis operon that usually produces dhATX as main toxic congener. These strains are capable of uncontrolled proliferation during warm seasons, during which their scums are a threat to mammals, especially dogs. We recommend that such blooms be reported and that specific management procedures be developed.

## CRediT

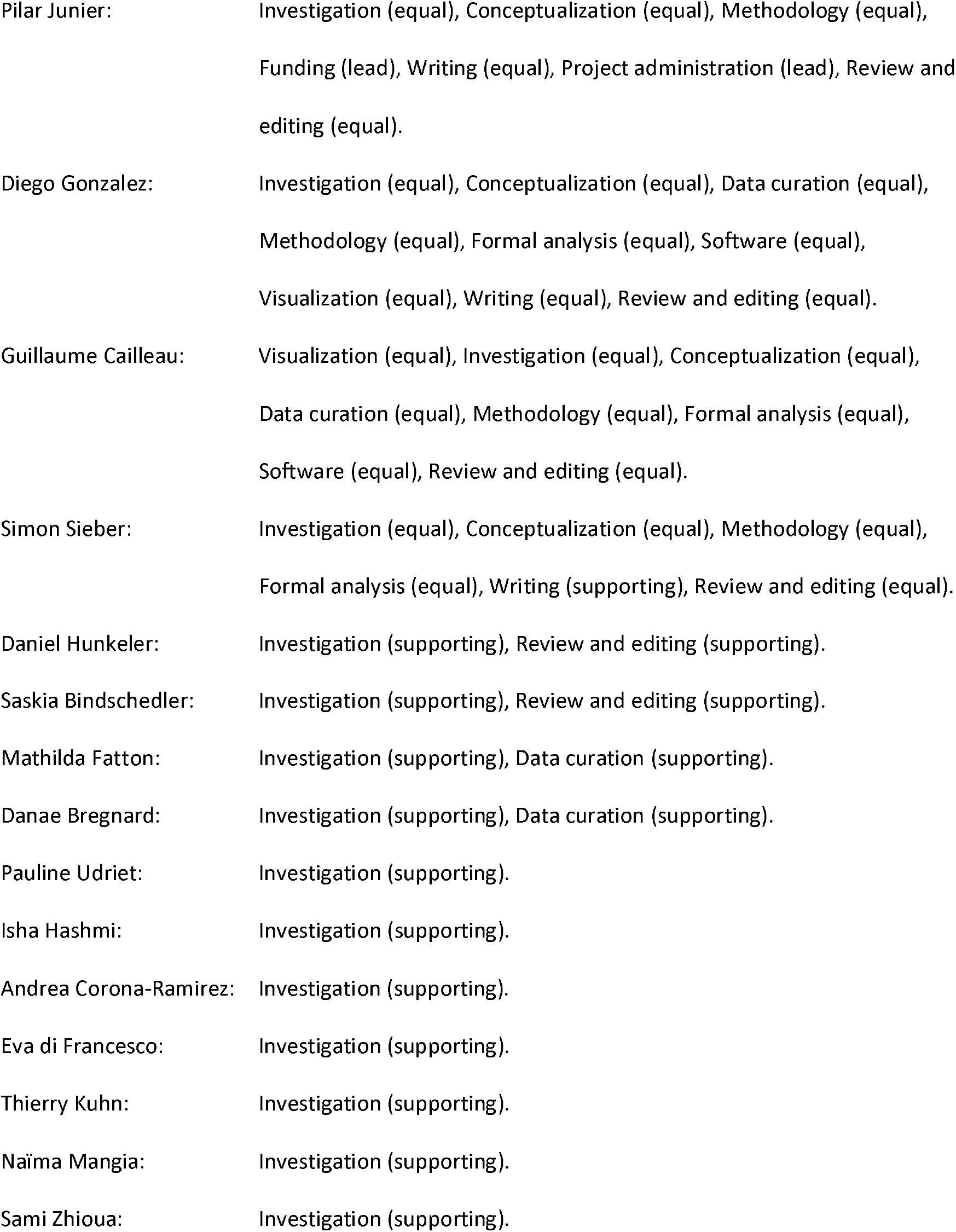

|  |  |
| --- | --- |
| Pilar Junier: | Investigation (equal), Conceptualization (equal), Methodology (equal),<br>Funding (lead), Writing (equal), Project administration (lead), Review and<br>editing (equal). |
| Diego Gonzalez: | Investigation (equal), Conceptualization (equal), Data curation (equal),<br>Methodology (equal), Formal analysis (equal), Software (equal),<br>Visualization (equal), Writing (equal), Review and editing (equal). |
| Guillaume Cailleau: | Visualization (equal), Investigation (equal), Conceptualization (equal),<br>Data curation (equal), Methodology (equal), Formal analysis (equal),<br>Software (equal), Review and editing (equal). |
| Simon Sieber: | Investigation (equal), Conceptualization (equal), Methodology (equal),<br>Formal analysis (equal), Writing (supporting), Review and editing (equal). |
| Daniel Hunkeler: | Investigation (supporting), Review and editing (supporting). |
| Saskia Bindschedler: | Investigation (supporting), Review and editing (supporting). |
| Mathilda Fatton: | Investigation (supporting), Data curation (supporting). |
| Danae Bregnard: | Investigation (supporting), Data curation (supporting). |
| Pauline Udriet: | Investigation (supporting). |
| Isha Hashmi: | Investigation (supporting). |
| Andrea Corona-Ramirez: | Investigation (supporting). |
| Eva di Francesco: | Investigation (supporting). |
| Thierry Kuhn: | Investigation (supporting). |
| Naïma Mangia: | Investigation (supporting). |
| Sami Zhioua: | Investigation (supporting). |

## Supporting information

STable 1

STable 2

STable 3

STable 4

## Acknowledgments

We thank Daniel Borel and Nathan Villat for their help during the sampling campaign, as well as Céline Terrettaz for follow up work. D.G’s work was supported by the *Swiss National Science Foundation* (PZ00P3_180142 to D.G., 2020-2023) and the *Velux Foundation* (grant 1814 to P.J. and D.G., 2023).

## Material and methods

### Sampling

The sampling sites corresponded to three areas: (1) the mouth of the Areuse river, just before it connects to Lake Neuchâtel; (2) River delta, and (3) by a dam on the river in Noiraigue. The two first samples were taken at the mouth on July 31st, 2020, right after the death of several dogs, and on August 16, 2020, as part of a first monitoring of the evolution of the proliferation. On September 8th, 2020, a sampling campaign took place to establish the extent of the zone where cyanobacterial mats developed in the river and in the lake. The first series of samples came from the river banks, mats or sediments on rocks and other emerged surfaces (23 samples; codes A to Y). In addition, samples from underwater mats and sediments from the river mouth and from the delta were obtained by diving (numerical codes below 100). Finally, on September 11, 2020, a second sampling took place near the town of Noiraigue and included three stations: directly in the town, near an abandoned dam and upstream of the current dam (7 samples; numerical codes 100 and above; Table S1). All samples were collected using a spatula to transfer cyanobacterial mats/biomass into tubes as free of contamination as possible.

### DNA extraction

500 mg of biomass were added to Lysing Matrix E tubes containing phosphate and MT buffer and vortexed. The tubes were then placed in a Tissue Lyser for 10 minutes at 50Hz, with intermittent cooling on ice. After centrifugation to remove debris, the supernatant was transferred to new tubes. For protein precipitation, Protein Precipitation Solution was added to the supernatant. The mixture was left to precipitate at 4°C for 10 minutes, followed by centrifugation. The Binding Matrix suspension was added to the supernatant. After incubation, the mixture was filtered through SPIN™ filters; multiple rounds of filtration were conducted if necessary. DNA was eluted from the matrix at 55°C in 100µl H2O. Storage was done at -20°C.

### 16S PCR for Cyanobacteria identification

A 40-cycle polymerase chain reaction (PCR) was performed using specific primers for cyanobacteria, Cya106f (5’-CGG ACG GGT GAG TAA CGC GTG A-3’) and Cya781R(b) (5’-GAC TAC AGG GGT ATC TAA TCC CCT T-3’) with the AllinTM Red Taq DNA Polymerase (highQu GmbH, Kraichtal, Germany). Annealing temperature was 58°C. The amplicon was sequenced using the Sanger method (Microsynth A.G., Switzerland). The amplified DNA was then purified by depositing the PCR products in a 96-well MultiScreen PCR plate (Millipore). A vacuum of 15 inHg was applied to the wells containing the PCR products until all the wells were dry (12-15 minutes). 30 µl of sterile ultrapure water was added to the wells, which were then allowed to stand for 1 min. The samples were resuspended in ultrapure water by back and forth pipetting and transferred to Eppendorfs with a volume of 0.5 ml. Purified PCR products were quantified again using the Qubit fluorometer before being sent to Fasteris for Sanger sequencing. Each PCR product was sequenced in both directions. After sequencing, each sample had a forward sequence and a reverse sequence. These sequences were then compared to a database available online (GenBank®) using the NCBI Basic Local Alignment Tool (BLAST) web interface ^48^.

### 16S community sequencing

The V3-V4 region (universal primers Bakt_341F 5’-CCT ACG GGN GGC WGC AG-3’ and Bakt_805R 5’-GAC TAC HVG GGT ATC TAA TCC-3’) ^49^ was amplified with a sample barcoding to allow multiplexing and adapter ligation to enable sequencing on the Illumina MiSeq platform (2 x 300 bp paired end reads). Demultiplexed and trimmed sequence reads provided by Fasteris were processed using *QIIME2* ^50^ with *dada2* ^51^ for the denoising step. Reads were truncated to 478 bases, the optimal length based of q-scores; a minimal overlap of at least 12 identical bases between pair-end reads was required to build a full sequence. The sequences were taxonomically classified using *QIIME2*’s *VSEARCH*-based consensus taxonomy classifier ^52^ with the *Silva* database ^53^, release 138, for 16S rRNA genes. The Silva database was fine-tuned using the RESCRIPt QIIME2 plugin ^54^, provided by the QIIME2 Team. Once the ASVs were determined, all ASVs represented in the template-free control samples were excluded from the corresponding culture collection. The co-occurrence network analysis was performed using *SpiecEasi* ^55^.

### Pacific Biosciences (Pacbio) SMRT shotgun metagenomics

1µg DNA extracted from four environmental samples (A, N, 600, X) was used to prepare standard bacterial Pacbio sequencing libraries (insert size: 9000 bps), Libraries were purified using Ampure beads and bound using Sequel® II Binding Kit 2.2. The libraries were sequenced on a Sequel II instrument as part of a 6-sample multiplexed run. Read count per sample was between 166’000 and 345’000, base count between 600M and 1800M. High fidelity reads were generated using Pacbio CCS.

### Genome assemblies

Genome assemblies were generated from high-fidelity reads only using *flye* (v. 2.9.2) with the --pacbio-hifi, --meta and --min-overlap 1000 options and three rounds of polishing; assemblies were further polished using *racon* (v. 1.5.0) with default parameters; *dnaA* (genome) and *parA* (plasmids) homologs were used to fix the start of circular genomes using *circlator* (v. 1.5.5). Contigs were assigned to taxonomical categories using *kraken2* (online version on the bv-brc.org website). Contigs larger than 20kbps were binned using *Rtsne* (pca=TRUE, perplexity=5, theta=0.1, check_duplicates = F) based on average observed over expected pentanucleotide frequency values calculated based on an in-house script. Coverage values were derived from *flye* assembly reports. Average Nucleotide identity was calculated using the *pyani* script (https://github.com/widdowquinn/pyani). 16S rRNA gene comparisons were done using the *pid* function (“PID2” method) of the *Biostrings* R package ^56^. Digital DNA-DNA hybridization was carried out using the GGDC 3.0 with Formula 2 (identities / HSP length) (https://ggdc.dsmz.de/) ^36^. D1-D1’ helix folding prediction was carried out using the *RNAfold* web server (ViennaRNA Package, v. 2.6.4) ^57^.

### Gene prediction and annotation

Standard annotation was carried out through the NCBI Prokaryotic Genome Annotation Pipeline (PGAP). Protein classification and pathway analyses were done using both the Kyoto Encyclopedia of Genes and Genomes (KEGG) online service (*blastKOALA*) and the comparative systems analysis of the BV-BRC website. The genomes used for comparisons were the following: *Oscillatoria_nigro-viridis* PCC_7112 (GCA_000317475.1), *Tychonema* sp. BBK16 (GCA_021648855.1), *Tychonema bourrellyi* FEM_GT703 (GCA_002412335.2), *Microcoleus* sp. CAWBG58 (GCA_020883045.1), *Microcoleus* sp. PH2017_15_JOR_U_A (GCA_020738445.1), *Microcoleus* sp. EPA2 (GCA_020882975.1), *Microcoleus* sp. CAWBG639 (GCA_020883025.1). For American *Microcoleus* strains PTRS1-3 (SRR10997082-4) and Canadian *Microcoleus* sequences (W3-F: SRR21374384, W3-E: SRR21374385, W5: SRR21374391, W3-J: SRR21374397), contigs were assembled from traces using *metaspades* (v. 3.15.5) with standard options. Proteome comparisons were carried out using the Proteome comparison service of BV-BRC (*Microcoleus* sp. strain A or *Oscillatoria nigro-viridis* PCC 7112 as reference proteomes) and plotted on the reference genome using the *circlize* R library. The secondary metabolite biosynthesis clusters were predicted using the online *antiSMASH* service.

### Phylogenetic trees

356 complete cyanobacterial genomes (nucleotides, proteomes, and annotations) were downloaded from NCBI. Large protein-based phylogenetic trees were built using *fasttree* with -wag and -gamma options from a multiple alignment, made using *muscle* or *kalign* with default parameters, of 28 concatenated conserved proteins (essentially ribosomal proteins). Large 16S rRNA phylogenetic trees were built using *fasttree* with -gtr option from a multiple alignment made using *muscle* with default parameters. A phylogenetic tree based on 28 concatenated conserved proteins for strains close to *Microcoleus anatoxicus* was built using *iqtree2* with automated optimization of the transition matrix; alignment was made using *muscle*.

### Data availability

Metagenome-assembled genome sequences were deposited on the NCBI repository (bioproject PRJNA943188). 16S rRNA data were deposited NCBI (bioproject XXXXX).

### Plots and figures

Figures were made using *R* (v. 3.14). The following *R* libraries were used: *ggplot2, pheatmap, circlize, biostrings, igraph*.

### Analysis of anatoxin-a and dihydroanatoxin-a

The sample collected in early August was analyzed by Ultra-high-performance liquid chromatography (UHPLC) coupled with a high-resolution mass spectrometer (HR-MS) [32]. The samples collected in September (STable 1) were analyzed using a UHPLC coupled with a triple quadrupole MS (TQMS) and with a selected reaction monitoring strategy for ATX as published previously [33]. Briefly, the cyanobacteria samples were centrifuged (30 min, 4000 g) and the supernatant was discarded. Water (30 mL) was added to the biomass, the mixture was vortexed and centrifuged (30 min, 4000 g), and the solution was decanted and discarded. An aqueous MeOH soln. (30 mL, 50%) was added to the biomass, the mixture was sonicated (3 x 3min), centrifuged (30 min, 4000 g), and the supernatant was collected and dried under a gentle flow of nitrogen. An aqueous MeOH soln. (50%) was added to the samples to afford a concentration of 1 mg/mL of extract and the samples were directly analyzed by UHPLC-ESI-MS in positive mode (Ultimate 3000 LC, Thermo Fisher Scientific, coupled to a TSQ Quantum Ultra, Thermo Fisher Scientific). The column used was the Kinetex Polar C18 (50 x 2.1 mm, 2.6 µm), the flow rate was set up to 0.4 ml/min, the column oven temperature was at 40°C, and the solvent system used was composed of A (H_2_O with 0.1% HCO_2_H) and B (MeCN with 0.1% HCO_2_H). The column was pre-equilibrated with 0% B for 1min, the gradient increased to 60% B over 3.5 min and to 100% over 0.05 min, and was kept to 100% B for 1.24 min. The detection was achieved using the SRM mode as previously reported ^33^ and a fragmentation from 166.2 to 130 was selected. For the quantification of dihydroanatoxin-a, the calibration curve was based on the SIM of anatoxin-a at 166.2 *m*/*z* and the concentration of dihydroanatoxin-a in the sample was calculated using the SIM mode at 168.2 *m*/*z*. The quantification was done using a calibration curve obtained using analytical standard solutions prepared with the commercially available anatoxin fumarate (Bio-Techne).

## Notes

### Competing Interest Statement

The authors have declared no competing interest.

### Summary of Updates

Recently, proliferations of benthic cyanobacteria producing derivatives of anatoxin-a have been reported in rivers all over the world. In three river systems, in New Zealand, the USA, and Canada, a cohesive cluster of Microcoleus strains was responsible for toxin production. Here, we document a similar toxigenic event that occurred at the mouth of the river Areuse in lake Neuchatel (Switzerland) and caused the death of several dogs. Using 16S RNA-based community analysis, we show that riverine benthic communities are dominated by Oscillatoriales and especially by Microcoleus strains. We correlate the detection of one sequence variant with the presence of anatoxin-a derivatives and use metagenomics to assemble a complete circular genome of the strain. The strain is distinct from the ones isolated in New Zealand, the USA, and Canada, but belongs to the same species; it shares significant traits with them, in particular a relatively small genome and incomplete vitamin biosynthetic pathways. Overall, our results suggest that the major anatoxin-a-associated benthic proliferations worldwide can be traced back to a single ubiquitous species, Microcoleus anatoxicus, rather than to a diversity of cyanobacterial lineages. We recommend that this species be monitored internationally in order to help predict and mitigate similar cyanotoxic events.

