## Supplementary figures for "A single *Microcoleus* species causes benthic cyanotoxic blooms worldwide"

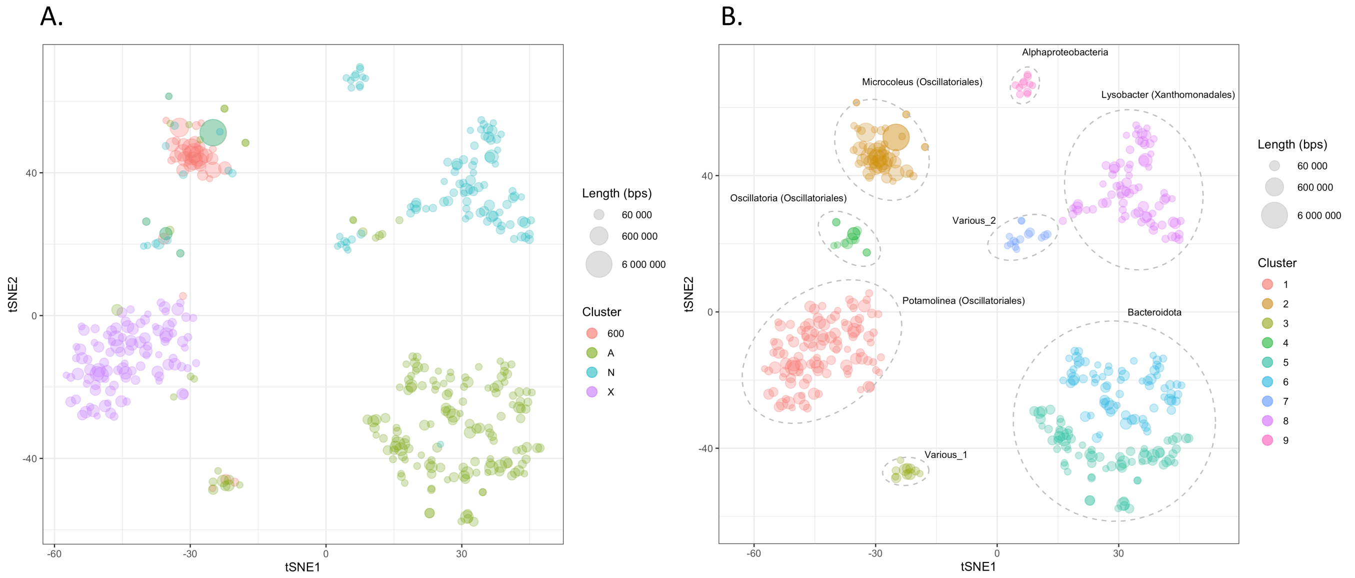


SFigure 1: tSNE clustering, based on pentanucleotide frequency, of large (>20kb) contigs from samples A, N, 600, and X. A. Mapping of the four different samples. B. Mapping and classification of the different clusters.

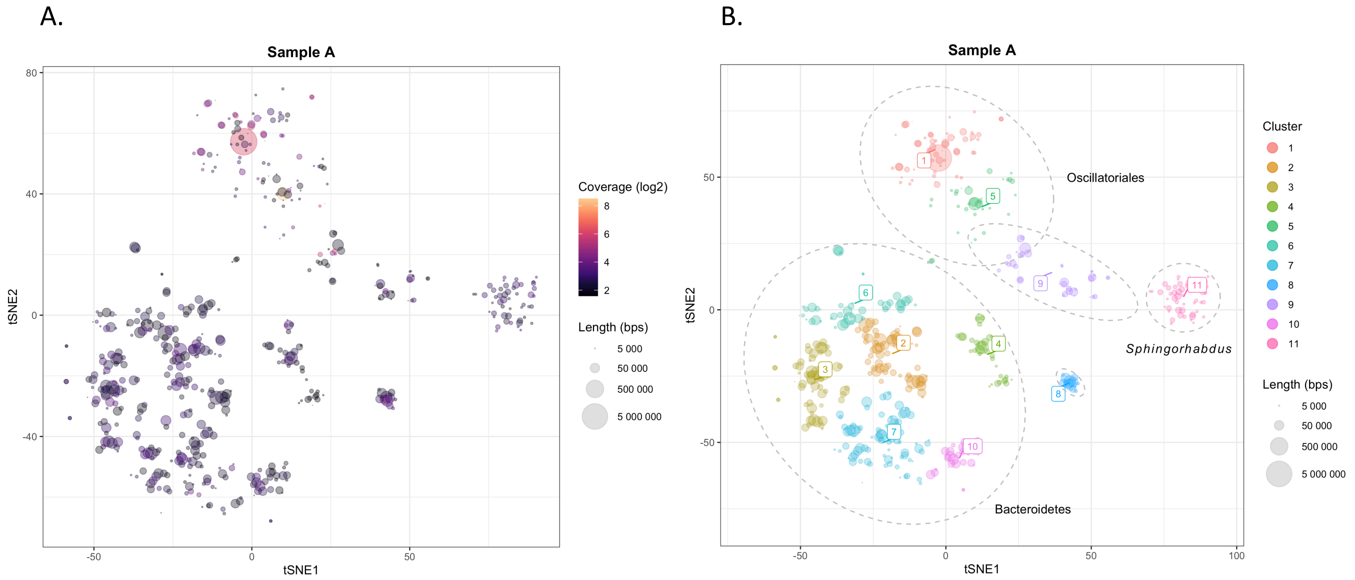


SFigure 2: tSNE clustering, based on pentanucleotide frequency, of large contigs from sample A. A. Mapping of the contig coverage values. B. Mapping and classification of the different clusters.

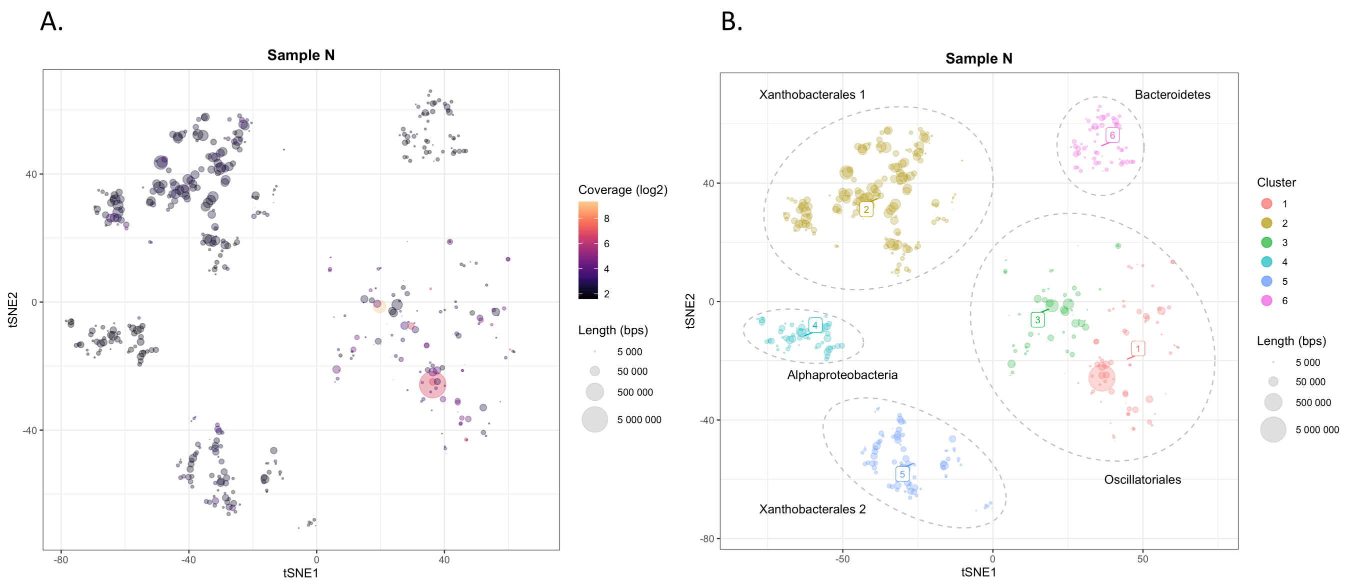


SFigure 3: tSNE clustering, based on pentanucleotide frequency, of contigs from sample N. A. Mapping of the contig coverage values. B. Mapping and classification of the different clusters.

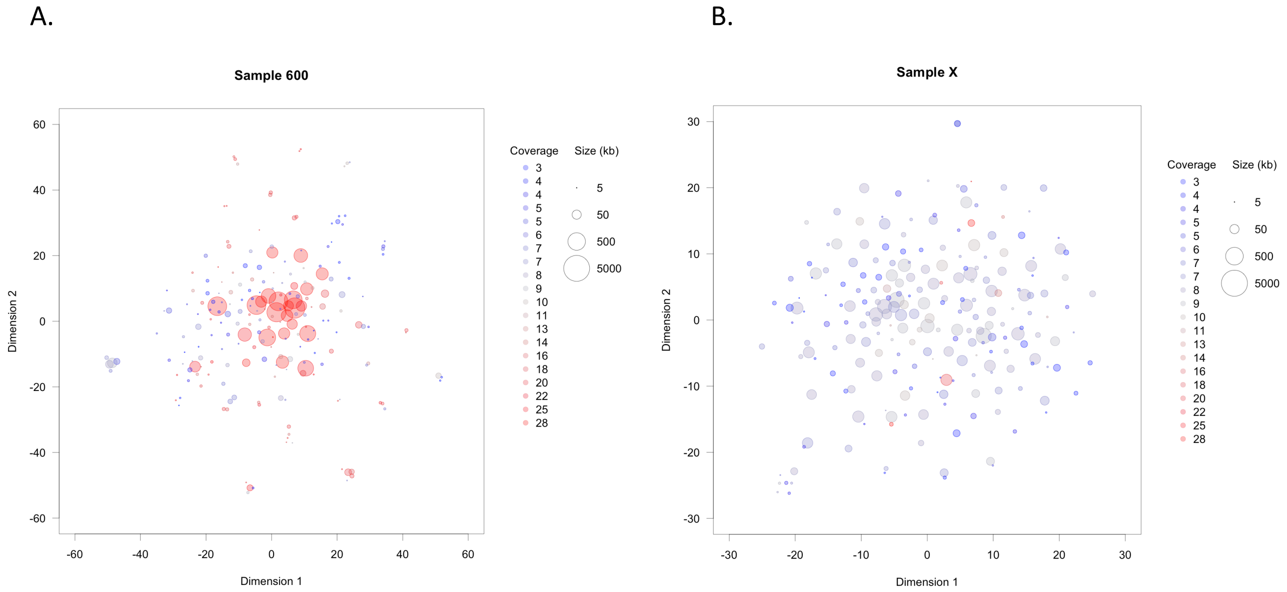


SFigure 4: tSNE clustering, based on pentanucleotide frequency, of contigs from sample 600 (A) et X (B). Contigs are almost all classified as Cyanobacteria.

A.

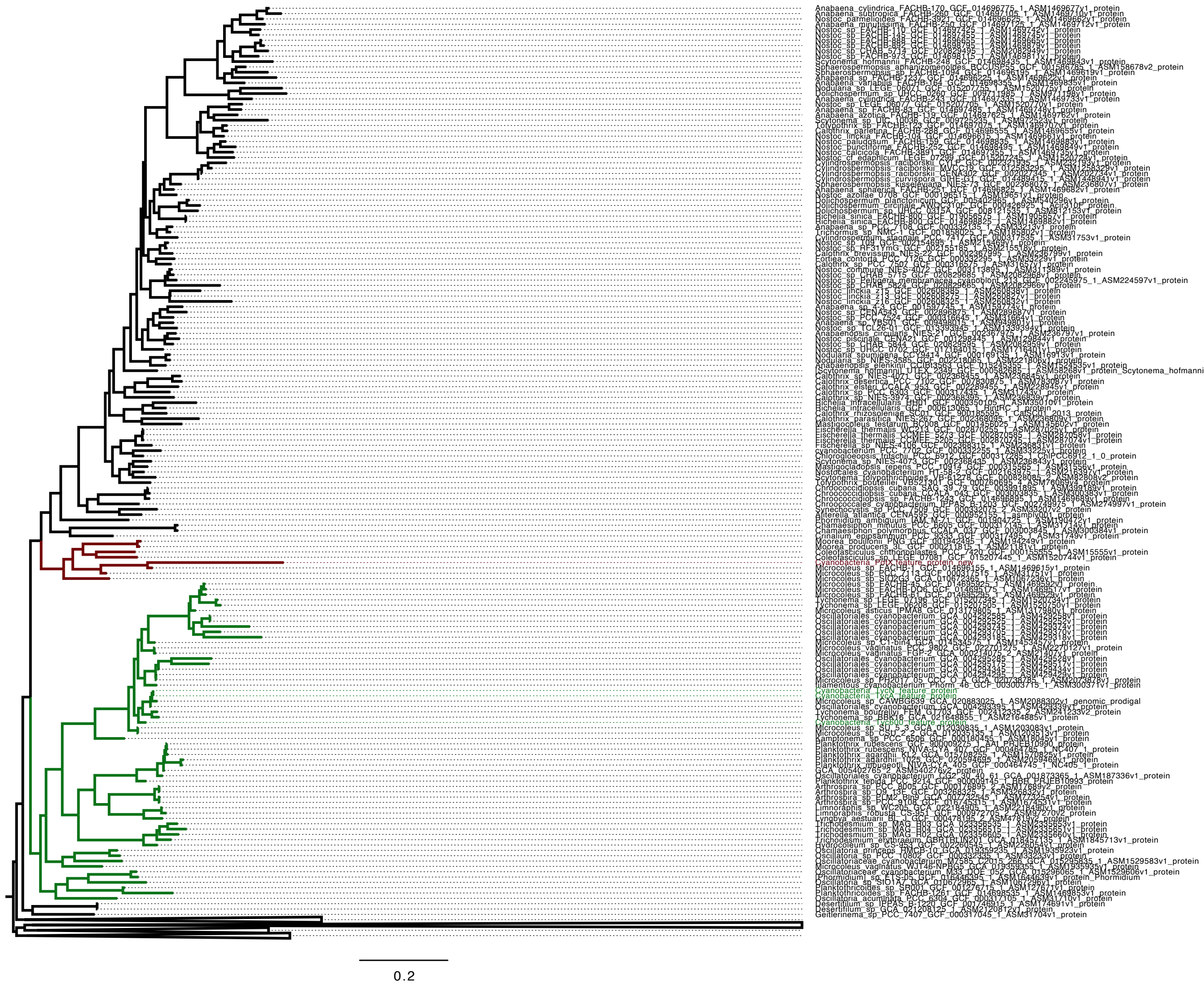

B.

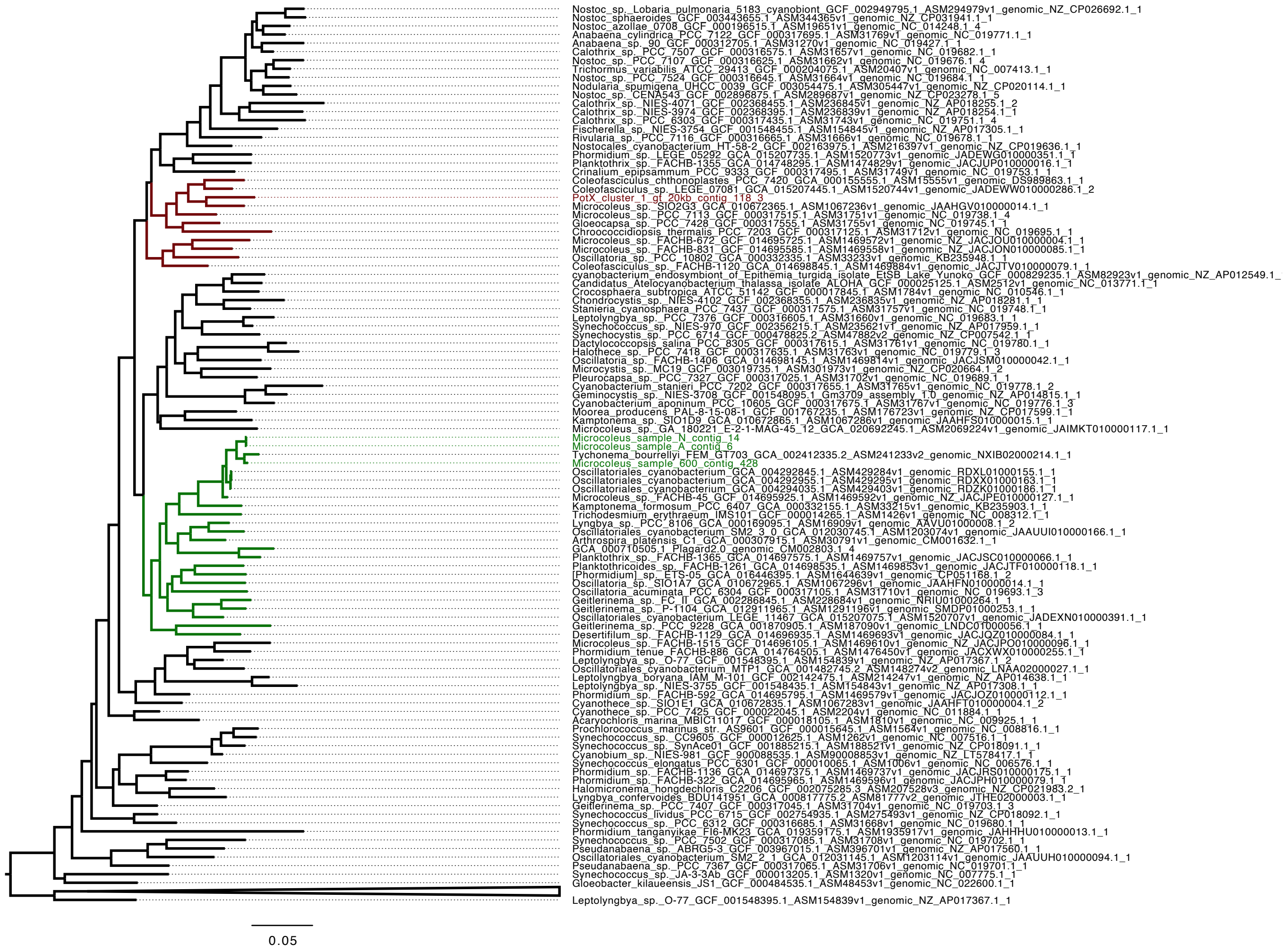


SFigure 5. Phylogenetic trees showing the position of the four genomes assembled from samples from the Areuse based on concatenated ribosomal proteins (A) or the 16S RNA sequence (B). In green, the *Microcoleus* genomes, in red the *Potamolinea* genome.

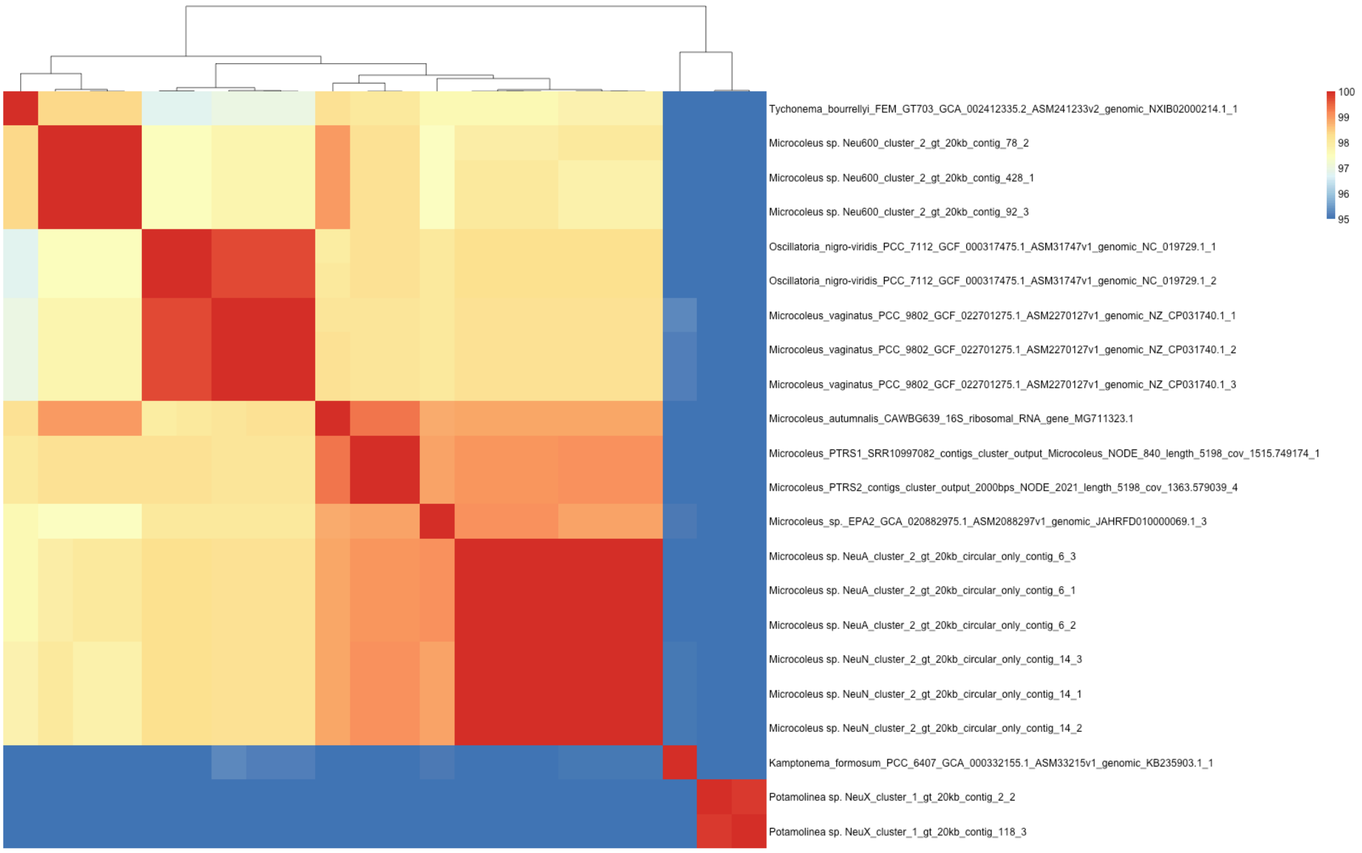


SFigure 6: Pairwise percent identity over the whole 16S rRNA gene between *Microcoleus*-related strains. Strains NeuA, NeuN, PTRS, EPA2, and CAWBG639 are all above 98% pairwise identity, which suggests that they belong to the same species. Values <95% were set at 95% to give a better color resolution to the higher percent identity range.

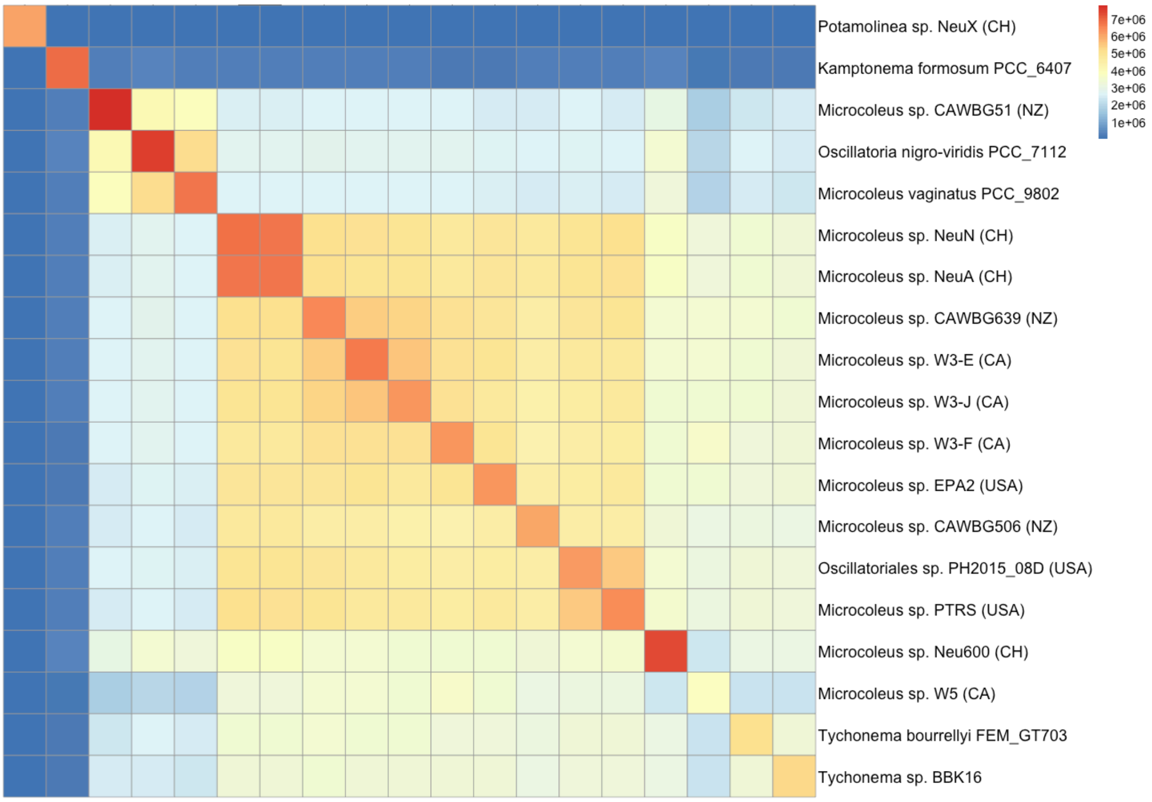


SFigure 7: Alignment lengths between representative genomes within the *Microcoleus-Tychonema* group.

| | | |
|---|---|---|
| 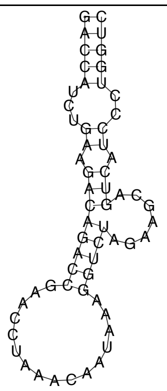 | 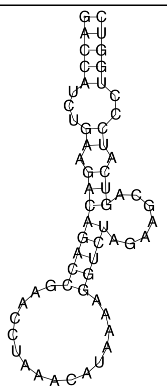 | 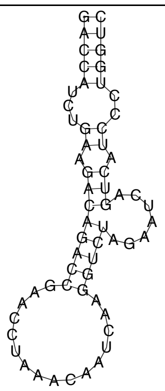 |
| *Microcoleus* sp. NeuA | *Microcoleus anatoxicus* strain PTRS-2 | *Microcoleus* sp. EPA2 |
| 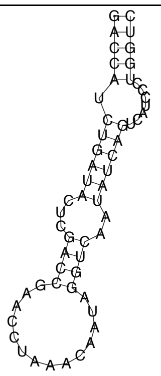 | 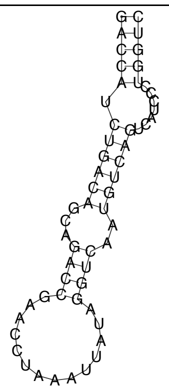 | 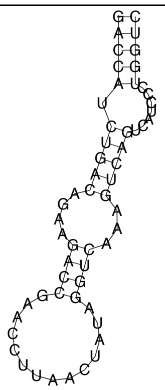 |
| *Microcoleus vaginatus* strain PCC_9802 | *Microcoleus* sp. Neu600 | *Tychonema bourrellyi* strain GT703 |

SFigure 8: Secondary structure of the of D1-D1' helices of the 16S-23S ribosomal RNA gene ITS region from selected genomes. *Microcoleus anatoxicus* strain PTRS2, Microcoleus sp. NeuA, and Microcoleus sp. EPA2 share a same D1-D1' helix secondary structure. Microcoleus sp. 600 shares a same structure with *Microcoleus vaginatus* strain PCC_9802.

A.

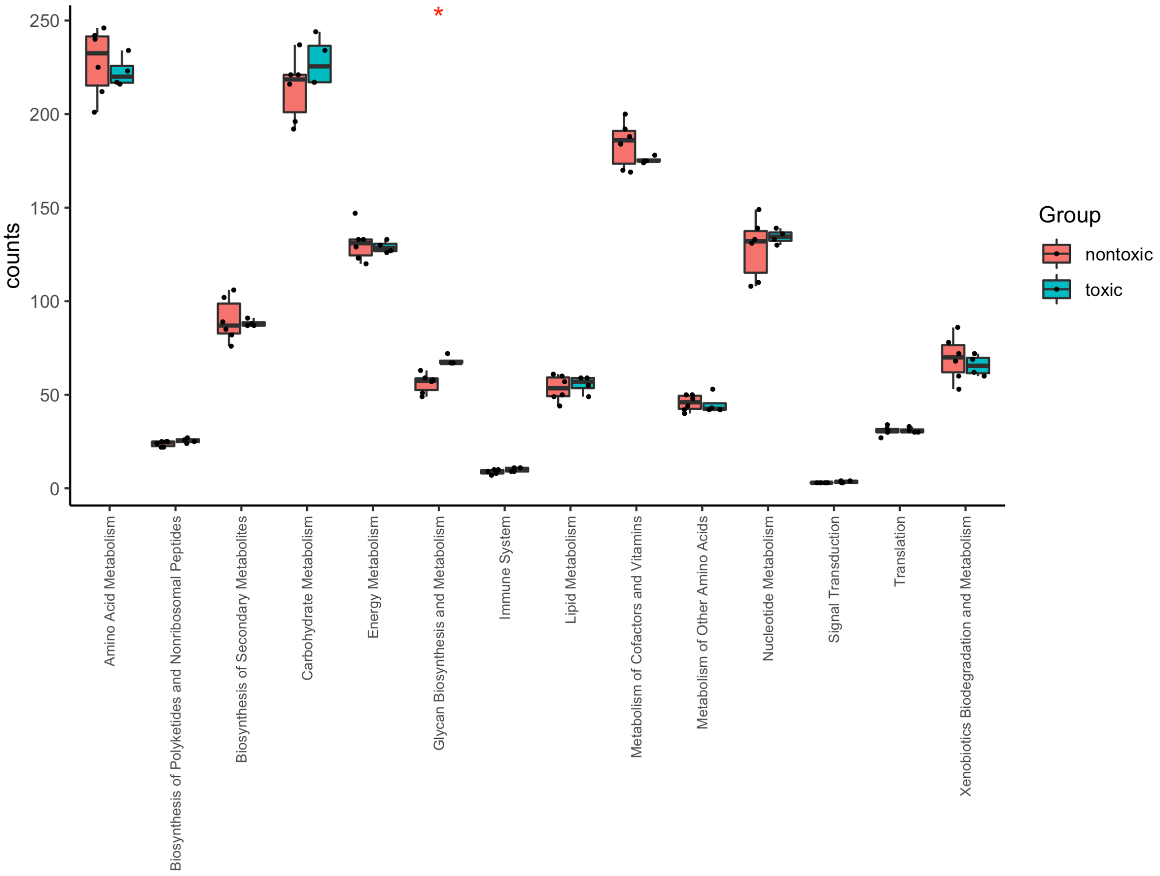


B.

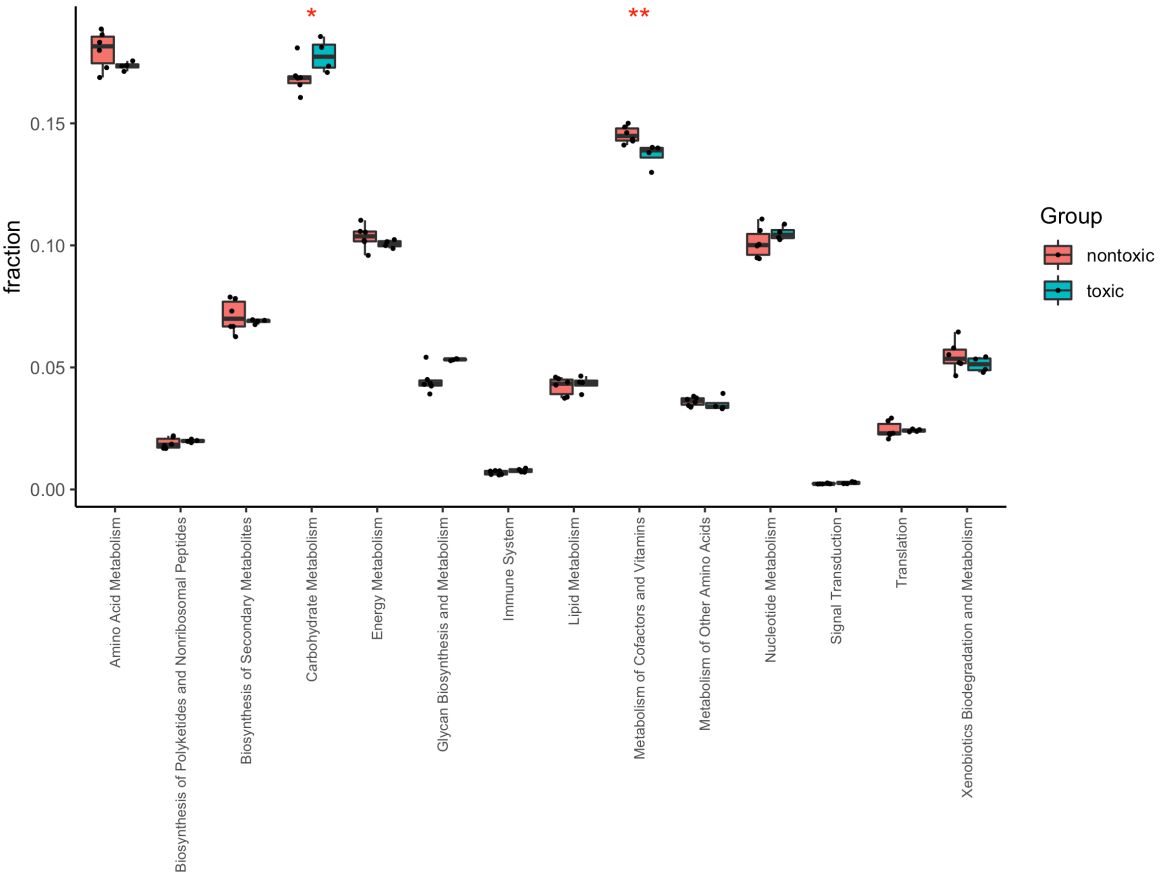


SFigure 9: Differences in protein function (main pathway) between *Microcoleus* strains that produce (toxic) or do not produce (nontoxic) anatoxin-a. A. Difference in absolute gene counts for a particular category. B. Difference in

the fraction of the genome for a particular category. List of toxic strains: *Microcoleus* sp. PRS1, CAWBG639, EPA2, NeuA; list of nontoxic strains: *Microcoleus* sp. CAWBG58, PH2017_15_JOR_U_A and Neu600, *Oscillatoria nigro-viridis* PCC 7112, *Tychonema* sp. FEM_GT703 and BBK16. Statistical test: unpaired two-samples Wilcoxon test, Holm correction for multiple testing. One red star indicates p-value<0.05; two stars indicate p-value<0.01. Functional annotation from PATRIC.

A.

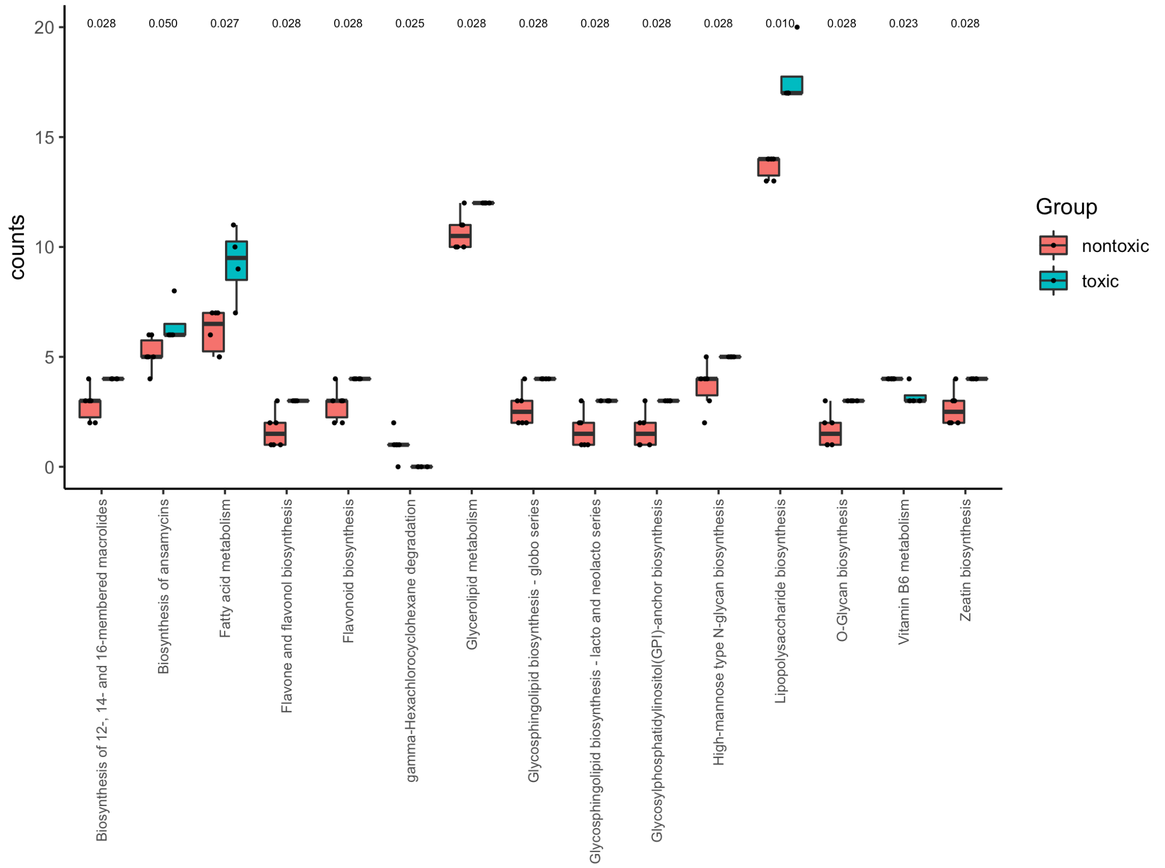


B.

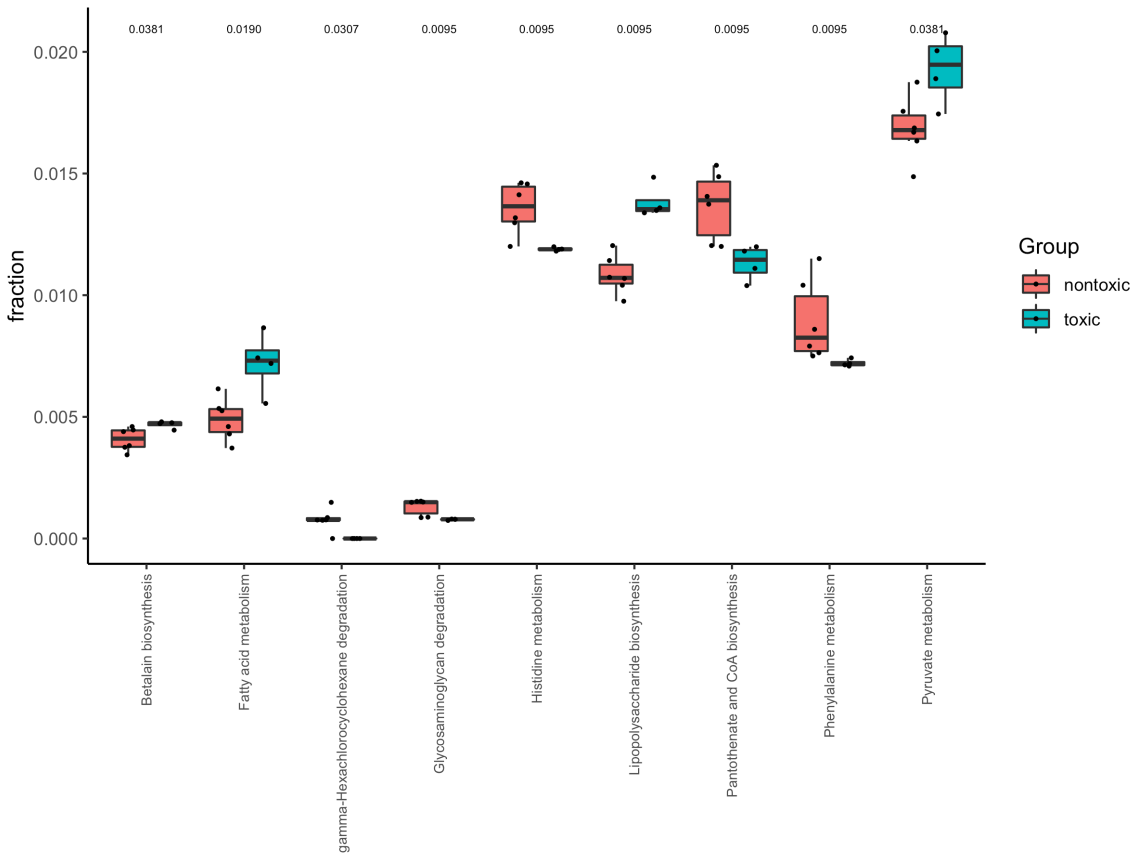


SFigure 10: Differences in protein function (subpathways) between *Microcoleus* strains that produce (toxic) or do not produce (nontoxic) anatoxin-a. A. Difference in absolute gene counts for a particular category. B. Difference in the fraction of the genome for a particular category. List of toxic strains: *Microcoleus* sp. PRS1, CAWBG639, EPA2,

NeuA; list of nontoxic strains: *Microcoleus* sp. CAWBG58, PH2017_15_JOR_U_A and Neu600, *Oscillatoria nigro-viridis* PCC 7112, *Tychonema* sp. FEM_GT703 and BBK16. Only statistically significant categories are shown (unpaired two-samples Wilcoxon test, Holm correction for multiple testing, pvalue<0.05); p-value is indicated above the boxes. Functional annotation from PATRIC.

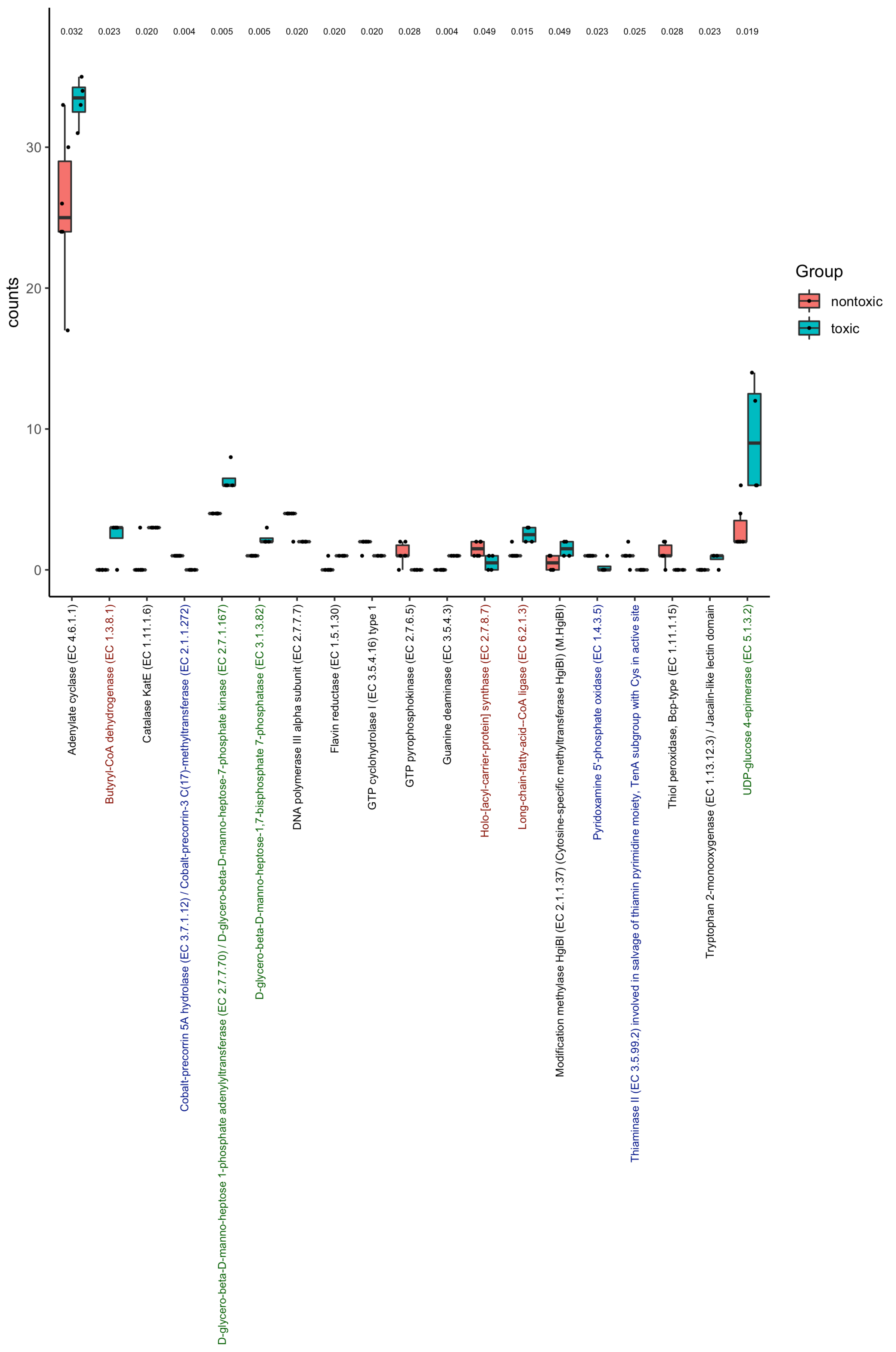


SFigure 11: Differences in protein function (subpathways) between *Microcoleus* strains that produce (toxic) or do not produce (nontoxic) anatoxin-a. List of toxic strains: *Microcoleus* sp. PRS1, CAWBG639, EPA2, NeuA; list of nontoxic strains: *Microcoleus* sp. CAWBG58, PH2017_15_JOR_U_A and Neu600, *Oscillatoria nigro-viridis* PCC 7112, *Tychonema* sp. FEM_GT703 and BBK16. Only statistically significant categories are shown (unpaired two-samples Wilcoxon test, Holm correction for multiple testing, pvalue<0.05); p-value is indicated above the boxes. Labels are colored in red for proteins involved in fatty acid metabolism, in blue for proteins involved in vitamin metabolism, and in green for proteins involved in sugar or polysaccharide metabolism. Functional annotation from PATRIC.
